# Brachyury expression levels predict lineage potential and axis-forming ability of *in vitro* derived neuromesodermal progenitors

**DOI:** 10.1101/2025.05.15.654369

**Authors:** Anahí Binagui-Casas, Anna Granés, Alberto Ceccarelli, Filip J. Wymeersch, Matthew French, Rosa Portero, Jen Annoh, Yali Huang, Eleni Karagianni, Frederick C.K. Wong, A. Sophie Brumm, Daniel Lopez Ramajo, Minoru Takasato, Sally Lowell, Osvaldo Chara, Valerie Wilson

## Abstract

Neuromesodermal progenitors (NMPs) produce the spinal cord and musculoskeleton in the elongating anterior-posterior axis. *In vivo*, NMPs possess dual potency, coinciding with regions coexpressing SOX2 and Brachyury (TBXT). *In vitro*, SOX2/TBXT co-expressing cells can be produced from pluripotent cells and, like their *in vivo* counterparts, can produce neural tube and somitic mesoderm. However, the functional characteristics of *in vitro* SOX2/TBXT co-expressing cells remain unclear, confounding comparisons with *in vivo* data. To address this, we developed a dual *Sox2/Tbxt* reporter mouse ESC line. SOX2/TBXT reporter-positive cells emerge *in vitro* from pluripotent populations with dynamics that mirror their appearance in the embryo. Purified SOX2/TBXT co-expressing populations can differentiate towards neurectoderm or mesoderm, including lateral mesoderm upon BMP stimulation. In gastruloids, quantitative live imaging shows that WNT or NOTCH inhibition rapidly leads to downregulation of TBXT expression and diminished axial extension. We show that clonally plated SOX2/TBXT co-expressing cells are bipotent NMPs that can also self-propagate. By combining clonal analysis with mathematical modelling, we identify two thresholds of SOX2/TBXT expression, switching clonal output from neural- to mesoderm-biased, and from mesoderm-biased to mesoderm-specified. Media and substrate composition alter the lineage outcomes of *in vitro* derived mouse NMPs. Thus, this *Sox2/Tbxt* double reporter cell line provides support for unsuspected heterogeneity in NMPs, together with evidence for a role of these transcription factors in directing cell fate to drive axis elongation.

## Introduction

In vertebrates, the head-to-tail (or anteroposterior, AP) body axis forms during early embryonic development. In mice, where this process is well characterised (Wymeersch et al., 2021), a transient population of progenitors, collectively called neuromesodermal competent cells (NMCs), produces both paraxial mesoderm and neurectoderm as the AP axis elongates in post-gastrulation stages. NMC populations across vertebrates harbour cells with functional heterogeneity, including *bona fide* bipotent neuromesodermal progenitors (NMPs) (Binagui-Casas et al., 2021). Importantly, the hallmark of NMPs is their dual fate towards both neuroectodermal and mesodermal lineages, two separate germ layers, at post-gastrulation stages (Guillot et al., 2021; Tzouanacou et al., 2009). Fate mapping has shown that cells with neuromesodermal fate are found at the posterior end of various vertebrate embryo species, and this region coincides with the co-expression of the transcription factors SOX2 and Brachyury (TBXT), reviewed in (Wymeersch et al., 2021).

Our current understanding of the literature suggests that NMPs in amniotes (chick, mouse) and anamniotes (fish, frog) are distinct. In amniotes, individual bi-potent NMPs are dual neural/mesodermal fated, while in anamniotes, neuromesodermal potency is maintained in NMC populations in the tail bud, but individual cells adopt either neural or mesodermal fates (Attardi et al., 2018; Davis & Kirschner, 2000; Guillot et al., 2021; Martin & Kimelman, 2012; Tzouanacou et al., 2009; Wymeersch et al., 2016). Thus, NMCs in the neuromesodermal-fated region in different species can consist of bona fide NMPs, or of a mixture of mono-fated cells of unknown potency.

*In vitro*, FGF and WNT treatment of pluripotent stem cells generates cells that resemble *in vivo* NMC populations both phenotypically and functionally: a high percentage of cells co-express SOX2/TBXT transcription factors, and these populations can differentiate into neural and mesodermal lineages (for a list of common protocols see Wymeersch et al., 2021). Indeed, at clonal level a proportion of *in vitro*-derived NMP-like cells expressing *Tbxt* (monitored with a fluorescent reporter) have dual potential for neural and mesodermal differentiation (Tsakiridis & Wilson, 2015). However, in these experiments, the knock-in reporter created a heterozygous null allele of TBXT (Fehling et al., 2003), a condition known to affect axial development (Gruneberg, 1958). Furthermore, the expression of SOX2 in the starting population was not measured, and therefore the relationship between SOX2 and TBXT coexpression, and bipotency is unknown.

Recent single cell RNA-seq studies have found that *in vitro* cultures harbouring NMC populations are transcriptionally heterogeneous, with varying levels of *Sox2* and *Tbxt*, as well as other NMC-specific transcripts such as *Msgn1*, *Nkx1-2,* and *Tbx6* (Dias et al., 2020; Edri, Hayward, Baillie-Johnson, et al., 2019; Edri, Hayward, Jawaid, et al., 2019; Gouti et al., 2017). One interpretation of this data is that *in vitro* NMC populations contain contaminating, non-NMP cells. Alternatively, the transcriptional heterogeneity in *Sox2* and *Tbxt* expressing cells themselves could reflect inherent functional heterogeneity at the single cell level given that transcription factors such as SOX2 elicit expression level-dependent function (Corsinotti et al., 2017), and that, in vivo, levels of TBXT and SOX2 protein appear to correlate with variations in NMC fate (French et al., 2025; Romanos et al., 2021; Wymeersch et al., 2016).

Addressing whether individual *in vitro* NMP-like cells co-expressing *Sox2* and *Tbxt* are bi-potent, and how heterogeneity in SOX2 and TBXT levels dictates differentiation outcomes requires the use of cells that report on the expression of both markers. Previous reporters have employed knock-in constructs that disrupt the function of these genes (Avilion et al., 2003; Fehling et al., 2003) or are additive bacterial artificial chromosome transgenics where there is a risk that expression does not accurately mirror endogenous expression (Koch et al., 2017).

Here we generated a minimally disruptive mouse embryonic stem cell (mESC) line with dual *Sox2/Tbxt* gene expression reporter activity and use it to examine live cell behaviours, population heterogeneity and functional potency of *in vitro* NMP-like cells. Clonal plating demonstrates that individual *Sox2/Tbxt* dual positive cells are *bona fide* bipotent progenitors, which can also self-propagate. Additionally, we show that rapid deceleration of axis elongation in gastruloids on WNT or NOTCH inhibition coincides with *Tbxt* downregulation, while changes in *Sox2* expression follow more slowly. We further uncover previously elusive lateral plate mesoderm potency within descendants of NMPs, an attribute shown only in bulk culture (Row et al., 2018).

Interestingly, we find that the *Sox2/Tbxt* co-expressing population is not functionally homogeneous: the levels of *Tbxt*-GFP expression predict clonal lineage bias towards mesoderm in individual progenitors. This intrinsic bias can be tuned by the culture conditions and cell substrates, mirroring the *in vivo* plasticity of NMC populations (Wymeersch et al., 2016).

## Results

### 1. A novel reporter line, STR-KI, accurately reports on SOX2 and TBXT expression and NMP differentiation trajectories *in vitro*

To characterise the dynamics of NMC emergence from mESCs, we constructed a 2A-peptide-linked dual reporter: SOX2::mCherry; TBXT::GFP, hereafter named SOX2-TBXT Reporter Knock-In (STR-KI). CRISPR knock-in transgenesis was used to create heterozygous knock-in alleles that replace the stop codon of the endogenous gene loci with T2A/P2A ‘self-cleaving’ (ribosome-skipping (Donnelly et al., 2001)) peptide, retaining the *Tbxt* and *Sox2* open reading frames in their entirety (Dewari et al., 2018). Since even partial loss of TBXT or SOX2 function affects differentiation (Avilion et al., 2003; Corsinotti et al., 2017; Gruneberg, 1958; Masui et al., 2007) we employed this strategy to minimise the chance of adverse phenotypic effects and maximise the faithfulness of the reporters (Fig. 1A). Chimeric embryos containing high contribution from STR-KI mESCs showed a normal phenotype, demonstrating that the gene targeting was indeed minimally disruptive to SOX2 and TBXT protein function. Furthermore, the fluorescent reporters in pre-streak and somite stage embryos recapitulated the *in vivo* expression pattern of TBXT and SOX2 in mesodermal and neural lineages, respectively (Fig. 1B, Suppl. Fig. 1C, D) (Wymeersch et al., 2016).In pre-streak chimeras, SOX2::mCherry (Sox2^mCh^ hereafter) was expressed throughout the epiblast, whereas TBXT::GFP (Tbxt^GFP^ hereafter) expression was undetectable (Suppl. Fig. 1C). By early somite stages onwards, Sox2^mCh^ was predominantly restricted to neural tissues, and to presumptive primordial germ cells at the base of the allantois, whereas Tbxt^GFP^-expressing cells were observed in the node, notochord and nascent mesoderm at the posterior end (Fig.1B and Suppl. 1D). Differentiation of STR-KI ESCs into NMP-like cells, following an established protocol (Gouti et al., 2014) produced mCherry and GFP double positive cells (Sox2^mCh^pos/Tbxt^GFP^pos) (Fig. 1C-F). Immunostaining and quantification of transcription factor (TF) intensity in these *in vitro* differentiated cells showed that the reporters faithfully co-localised with endogenous SOX2 and TBXT protein (Fig. 1C, D). For both reporters, we also observed a positive correlation between the levels of reporter and endogenous protein (Fig. 1C). We therefore adopt the working hypothesis that GFP and mCherry levels reflect TBXT and SOX2 protein production respectively, although differential posttranscriptional modifications affecting protein levels cannot be excluded.

**Figure 1.**
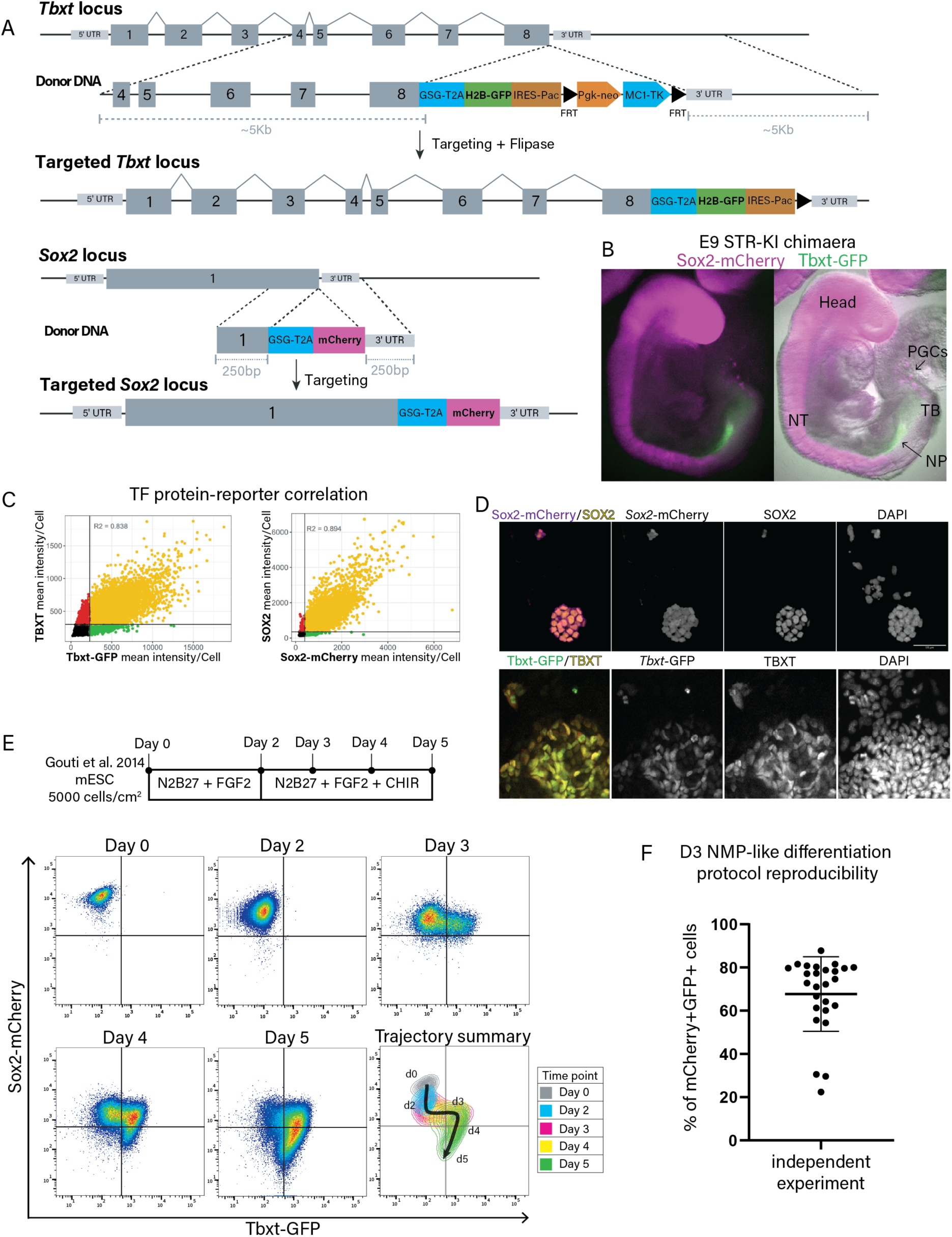
STR-KI accurately reports on SOX2 and TBXT expression and NMP differentiation trajectories *in vitro*. **A.** Schematic illustration of the targeting strategy to knock-in GFP and mCherry in the *Tbxt* and *Sox2* loci, respectively. **B.** Embryonic day 9 (E9) STR-KI chimaera with high STR-KI contribution shows appropriate localisation of fluorescent reporter expression: high mCherry (magenta) in the prospective brain, the neural tube (NT) and in primordial germ cells (PGCs). GFP (green) expression marks the notochordal plate (NP) and tail bud (TB). **C.** Correlation between immunofluorescent staining for SOX2 or TBXT, and their respective reporter protein in individual cells derived from *in vitro* NMP differentiation assays. **D.** Immunofluorescence images of differentiated STR-KI colonies following NMP-like differentiation (Gouti et al., 2014). Scale bar = 50μm. **E.** Representative example from a flow cytometry time-course analysis of STR-KI cells along the differentiation protocol shown in D, from mESCs (day 0) via NMP-like intermediate (at day 3) to day 5 (n=4). A trajectory summary highlights population dynamics. **F.** Quantification of the reproducibility of STR-KI mESC differentiation to day 3 NMP-like cells shows the percentages of Sox2^mCh^/Tbxt^GFP^double positive cells for a cohort of independent experiments (n=23; Mean ± SD: 67.42± 18.28%).

We next tracked the reporters’ dynamics over a time course of NMP differentiation (Fig. 1E). Briefly, this entails differentiation of ESC to an epiblast-like state, followed by treatment with the WNT agonist CHIR99021 (CHIR) and FGF2 from day (d) 2 onwards (Gouti et al., 2014). Cells appeared to move along a differentiation trajectory from Sox2^mCh^high pluripotent ESC to a Sox2^mCh^low, putative epiblast state, followed by appearance of Sox2^mCh^pos/Tbxt^GFP^pos (double positive cells) within 3 days (d). D3 differentiated cells exhibited a spectrum of Tbxt^GFP^ levels before exiting the double positive state towards Sox2^mCh^ negative (neg)/Tbxt^GFP^high mesoderm from d4. Consistent with previous reports, STR-KI ESC differentiation reproducibly resulted in the production of putative NMCs co-expressing mCherry and GFP at a rate of ∼67% of the population by d3 (Fig. 1F).

### 2. Expression dynamics of *Sox2* and *Tbxt in vitro* matches NMP emergence *in vivo*

A previous report demonstrated that *Sox2* expression in the caudal lateral epiblast is activated *de novo* in mouse via its N1 enhancer activity (Takemoto et al., 2011), corresponding approximately to the time at which NMCs are a detectable cell population (Wymeersch et al., 2016). Superficially, this would suggest that *Sox2* is secondarily activated in the *Tbxt*-expressing caudal lateral epiblast, contrasting with our observation of the acquisition of *Tbxt* in *Sox2*-expressing cells *in vitro* (Figure 1E). We therefore examined the origin of NMPs in the epiblast.

For this, we first analysed SOX2 and TBXT immunofluorescence in embryos staged between mid-streak and late headfold (Lawson & Wilson, 2025) (Figure 2A), including the neural plate (also known as pre-headfold) stage at which TBXT+SOX2+ putative NMPs emerge (Wymeersch et al., 2016). Up to early neural plate stage, SOX2 single positive cells were located anterior to the node, while cells posterior to the node expressed only TBXT. Double positive cells emerged at the node and just posteriorly to it during neural plate stage and constituted most of the cells at the posterior node from late neural plate stage onwards. Comparison of this staining pattern with focal electroporations at neural plate stage (n=2 anterior to node, n=3 posterior to node, Figure 2B, C) and previously published *in vivo* single cell clonal lineage tracing around the node (Forlani et al., 2003) (Figure 2C) shows that before neural plate stages, cells with dual neuromesodermal fate are located anterior to the node, while those posterior to it are exclusively mesoderm-fated. Thus, NMPs in the embryo emerge from the SOX2+ cell population anteriorly to the node and subsequently acquire TBXT expression (Figure 2D). This indicates that the acquisition of TBXT expression by SOX2+ cells in NMP differentiation conditions i*n vitro* mimics the *in vivo* ontogeny of NMPs.

**Figure 2.**
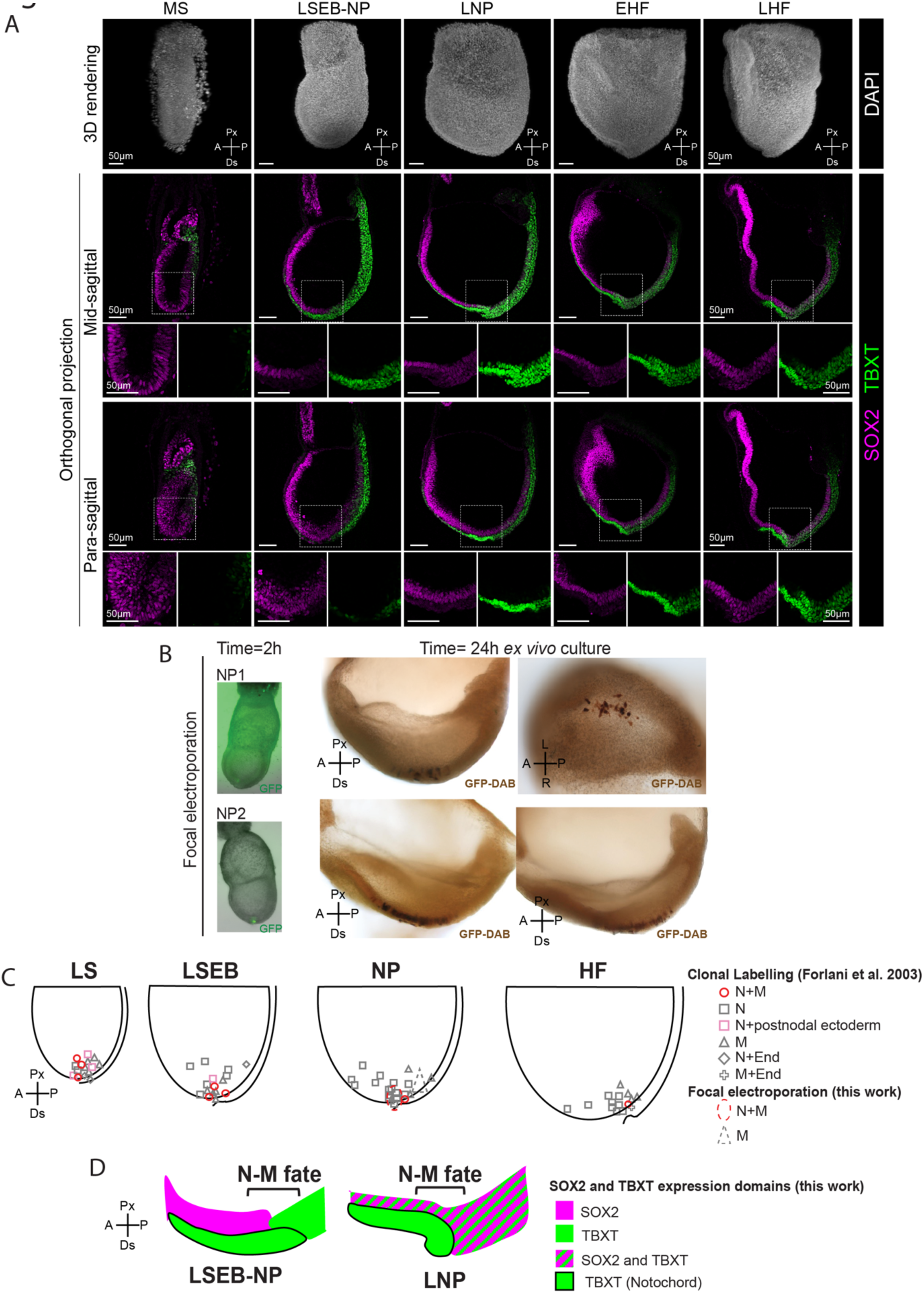
NMPs emerge *in vivo* from a pre-existing SOX2+TBXT-population. **A**. Immunofluorescent detection of SOX2 and TBXT in the fate-mapped region (boxed) over developmental time. Upper row: DAPI-stained 3D renderings of mouse embryos (dissected on embryonic days E6.5 (midstreak, MS), E7.0 (late streak early bud, LSEB), E7.5 (late neural plate, LNP) and E8.0 (early and late head fold, EHF, LHF); (Lawson & Wilson, 2025)). Lower rows: Orthogonal projections of wholemount immunostained mouse embryos for SOX2 (magenta) and TBXT (green), showing mid- and para-sagittal optical sections of the node area (projections represent 8 µm across the Z axis). All scale bars are 50 µm. **B**. Fate mapping at neural plate (NP) stage via focal electroporation of a ubiquitously-expressed CAG-GFP transgene into cells anterior to the node tracks descendants into the neural plate and anterior caudal lateral epiblast (n=2). Equivalent electroporation in the primitive streak (n=3) produces descendants in the mesoderm (data summarised in C). **C**. Focal electroporations superimposed on data from Forlani et al. 2003, where cells in or near the node at late streak (LS) to head fold (HF) stages were prospectively mapped to segments of the axis. Data has been re-displayed to indicate clones producing descendants in neurectoderm and/or mesoderm. **D**. Summary scheme of data in A-C. Tracing of LSEB-NP and NP stage node region showing expression of SOX2 and TBXT in epiblast and nascent notochord. Expression is non-overlapping at the midline in LSEB-NP transition stage embryo, while in LNP stage embryo onwards, SOX2+TBXT+ cells are evident. The anteroposterior extent of NM fate in the epiblast, extracted from diagrammatic representations in Forlani et al. at each stage is indicated, showing N-M fate is initially localized in the SOX2+ dorsal node region, which later becomes SOX2+TBXT+. Px, proximal; Ds, distal; A, anterior; P, posterior; L, left; R, right.

### 3. *In vitro* derived Sox2^mCh^/Tbxt^GFP^ co-expressing cells are NMCs

To characterise the cell populations emerging along the ESC-to-NMP-like cell trajectory, we probed the gene expression profiles of various subpopulations of diagnostic marker genes from d2 to d4 via qRT-PCR (Fig. 3A, B; Suppl. 2 A, B). Subpopulations were delineated based on the intensities of mCherry and GFP reporter detection in FACS. Comparison of mRNA expression levels of the endogenous *Sox2* and *Tbxt* with their respective reporters *mCherry* and *GFP* showed that in all sorted populations, mRNA expression reflected endogenous gene expression levels. The levels of *Sox2* and *Tbxt* mRNAs exceeded those of the corresponding reporters, as expected, since the reporter allele, which transcribes both endogenous mRNA and reporter, is heterozygous (Fig 3B). Interestingly, the putative NMC population (Sox2^mCh^pos/Tbxt^GFP^pos) displayed relatively constant Sox2^mCh^ fluorescence but a spectrum of Tbxt^GFP^ expression. Since the levels of Tbxt^GFP^ in the double-positive population increased before the appearance of Tbxt^GFP^ single positive mesoderm cells, we hypothesised that low and high Tbxt^GFP^ populations might represent different states in the wider NMC population. Specifically, high Tbxt^GFP^ positive cells might be further along a mesoderm differentiation trajectory compared to Tbxt^GFP^ low cells(French et al., 2025; Romanos et al., 2021; Wymeersch et al., 2016)(French et al., 2025; Romanos et al., 2021; Wymeersch et al., 2016)1).

**Figure 3.**
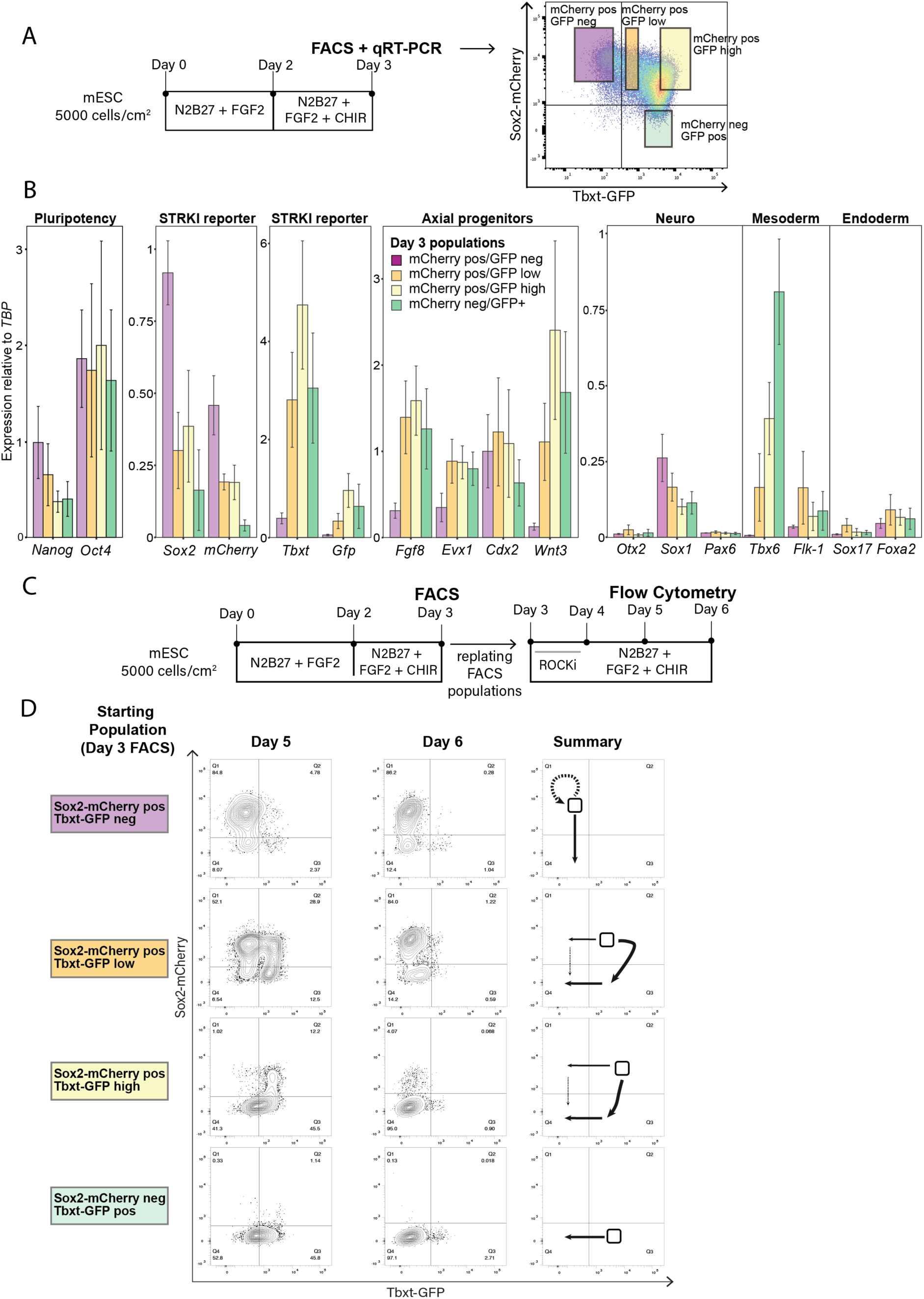
Gene expression and lineage potential analysis of Sox2^mCh^/Tbxt^GFP^ fluorescent subpopulations during differentiation. **A.** Differentiation scheme and representative gating strategy for FACS and further qRT-PCR analyses performed at day 3 of the differentiation protocol. **B**. qRT-PCR gene expression analysis of the populations in A. n=5 independent differentiation experiments. **C.** Post-d3 differentiation scheme in which populations in A were isolated by FACS, replated and further differentiated for an additional 3 days. **D.** Representative flow cytometry graphs of one time-course experiment (n=3, see Suppl. Fig. 2C), following the Sox2^mCh^ and Tbxt^GFP^ expression dynamics over three more days. A summary of the lineage outcomes in three experiments over time is provided. The thickness of arrows represents the relative contribution to the indicated trajectory, with dashed arrows indicating a presumed trajectory based on the proportions of cells at the end of the experiment.

To test this, we divided the NMC-like population into low-TBXT (Sox2^mCh^pos/Tbxt^GFP^low) and high-TBXT (Sox2^mCh^pos/Tbxt^GFP^high) expressing subpopulations. We analysed subpopulations based on marker gene expression: pluripotency-associated *Nanog* and *Oct4*; axial progenitor *Fgf8*, *Evx1*, *Cdx2* and *Wnt3a*; paraxial mesoderm *Tbx6;* lateral plate mesoderm (LPM) *Flk-1*; and neural-associated *Otx2, Sox1* and *Pax6*. Although both low-TBXT and high-TBXT populations expressed similar levels of NMC markers, high-TBXT cells were enriched for mesodermal markers such as *Wnt3a* and *Tbx6*, while expression of neural genes like *Sox1* were low. The absence of *Sox2*, coupled with high expression of *Tbx6* and *Flk-1* in both Sox2^mCh^neg/Tbxt^GFP^pos and Sox2^mCh^neg/Tbxt^GFP^neg (double negative) populations, together with the downregulation of NMC/caudal markers like *Fgf8* and *Wnt3a* from d3 to d4, suggests that these two populations represent nascent and differentiated mesoderm, respectively (Suppl. Fig. 2A, B). Finally, the expression of endoderm markers *Sox17* and *Foxa2* was universally low, suggesting negligible fractions of contaminant non-NMC cells. Thus, gene expression analysis in populations undergoing differentiation confirms that our double reporter cell line faithfully recapitulates endogenous gene expression and can be used to purify neural, mesodermal and NMC-like cell populations based on Sox2^mCh^ and Tbxt^GFP^ expression levels (Fig. 3B, Suppl. 2A, B). Furthermore, this analysis supports the idea that high levels of Tbxt^GFP^ mark a more mature mesodermal state than low Tbxt^GFP^ levels.

To test whether the apparently dynamic changes in Sox2^mCh^ and Tbxt^GFP^ cell populations during NMP differentiation (Fig. 1E) represent one or more lineage trajectories, the lineage potentials of sorted subpopulations at d3 of NMP differentiation were assayed by FACS over 3 further days in continuous FGF and CHIR, which are permissive for both neural and mesodermal differentiation (Gouti et al., 2014; Tsakiridis & Wilson, 2015) (Fig 3C, D Suppl. Fig. 2C). Most (>80%) Sox2^mCh^ single positive cells remained single positive. Together with the enrichment of *Sox1* and *Nanog* in the starting population (Fig 3B), this suggests a neural and/or epiblast identity of Sox2^mCh^ single positive cells (Wong et al., 2025). A small proportion of the progeny lost expression of both reporters, likely representing further maturation, since SOX2 is downregulated during neural differentiation . Sox2^mCh^pos/Tbxt^GFP^low cells also produced Sox2^mCh^ single positive cells by d6, but also a recapitulation of the pattern shown in Fig. 1E: emergence of Sox2^mCh^pos/Tbxt^GFP^high cells, followed by a high proportion of Sox2^mCh^neg/Tbxt^GFP^pos and later double negative cells. This suggests that Sox2^mCh^pos/Tbxt^GFP^low NMCs produce both neural and mesodermal cells. Interestingly, Sox2^mCh^pos/Tbxt^GFP^high cells produced few Sox2^mCh^ single positive cells and followed a predominantly mesoderm trajectory, producing Sox2^mCh^neg/Tbxt^GFP^pos and then double negative cells. Finally, Sox2^mCh^neg/Tbxt^GFP^pos populations only produced double negative cells. Together with its enrichment in expression of mesoderm markers, it suggests this is a mesoderm-committed population. Thus, using our dual reporter, we describe a lineage trajectory in which a largely self-sustaining SOX2-positive population generates a SOX2-positive;TBXT-low dual-potency population, which progresses through a mesoderm-biased SOX2-positive;TBXT-high state towards a nascent mesodermal TBXT-high and finally a double-negative state (Fig 3D, see summary). This indicates that the TBXT-high NMC population is predisposed towards mesoderm differentiation. Thus, within the Sox2^mCh^pos/Tbxt^GFP^pos double positive NMC population, there are apparent cells with differing competence to differentiate to neural or mesodermal outcomes.

Previous reports have shown that treatment of pluripotent cells with FGF2 and CHIR leads to the expression of both paraxial and lateral plate mesoderm (LPM) markers (Edri, Hayward, Baillie-Johnson, et al., 2019; Gouti et al., 2017). *In vivo*, while homotopic grafts of NMCs overwhelmingly give rise to paraxial mesoderm (Cambray & Wilson, 2007; Wymeersch et al., 2016), heterotopic grafts to the posterior primitive streak leads to LPM differentiation (Row et al., 2018). This suggests that NMPs may have LPM potency, although it cannot be excluded that this LPM arose from non-NMCs within the graft. To directly test the LPM-potency of NMCs, we examined the potential of FACS-enriched Sox2^mCh^pos /Tbxt^GFP^high cells to produce LPM lineages upon BMP treatment, which induces LPM differentiation in bulk NMP cultures (Row et al., 2018). We observed not only FLK-1 protein (a lateral plate/endothelial cell marker) labelling in colonies but also the formation of primitive tubes similar to those seen in classical endothelial assays (Taoudi et al., 2005) (Suppl. Fig. 2D, E). These findings indicate that Sox2^mCh^pos /Tbxt^GFP^high NMP-like cells can produce LPM. Thus, the SOX2-pos;TBXT-high mesoderm-fated population shows broad mesodermal potency, supporting the idea that *in vivo*, it is NMCs themselves, and not contaminating populations, that can contribute to both paraxial and lateral mesoderm.

### 4. Live imaging of STR-KI cells uncovers a role for TBXT in WNT and NOTCH-driven axis elongation

In vivo, wildtype TBXT levels are not only critical for NMP differentiation, but also to maintain axial elongation (Wymeersch et al., 2021). However, the dynamic relationship between TBXT, SOX2 and axial elongation remains untested. To determine in real time the relationship between reporter expression, axial elongation and factors known to affect both reporter expression and axial elongation, we next used live-tracking of STR-KI reporter expression. For this, we tracked Sox2^mCh^ and Tbxt^GFP^ expression *in vitro* in a standard gastruloid model of axis elongation, (Baillie-Johnson et al., 2015; van den Brink et al., 2014), in which aggregates of mESCs grown in N2B27 remain SOX2 pos/TBXT negative until 48 hours post aggregation (hpa), then exit pluripotency. At 48 hours, a 24-hour pulse of CHIR is used to activate WNT globally and ensure robust symmetry breaking. WNT activation transiently induces TBXT expression in all cells (van den Brink et al., 2014), before it resolves into polarized expression at the posterior pole by 72 hpa and is then followed by elongation. We first compared the dynamics of mesoderm differentiation in STR-KI-gastruloids with previous reports (Suppl. Fig. 2F). After treatment with CHIR, most cells became Tbxt^GFP^pos by 72 hpa, and down-regulated Sox2^mCh^. In the absence of CHIR, STR-KI cells remained Sox2^mCh^high, but a small proportion of cells began to express Tbxt^GFP^. These trends were maintained at the population level by 96 hpa (Suppl. Fig. 2F). We also observed downregulation of Sox2^mCh^, symmetry breaking and polarisation of Tbxt^GFP^ to the posterior growing axis only upon the CHIR pulse (Suppl. Fig. 2G). We conclude that our STR-KI reporter can generate gastruloids similar to other lines and thus it is ideally suited for studying TF dynamics within them.

We next assessed how the manipulation of WNT and NOTCH signalling pathways, which regulate mesoderm differentiation from NMPs (Cooper et al., 2024; Wymeersch et al., 2016), impact gastruloid axial elongation. We treated elongating gastruloids (96 hpa) with either WNT or NOTCH inhibitors (IWP-2 and LY411575 (LY), respectively) and live imaged them every hour for 24 hours to monitor major axis length and reporter expression (Fig. 4A, B). Both WNT and NOTCH inhibition decelerated major axis growth (elongation) starting from as early as 2-3h post treatment (Fig. 4C).

**Figure 4.**
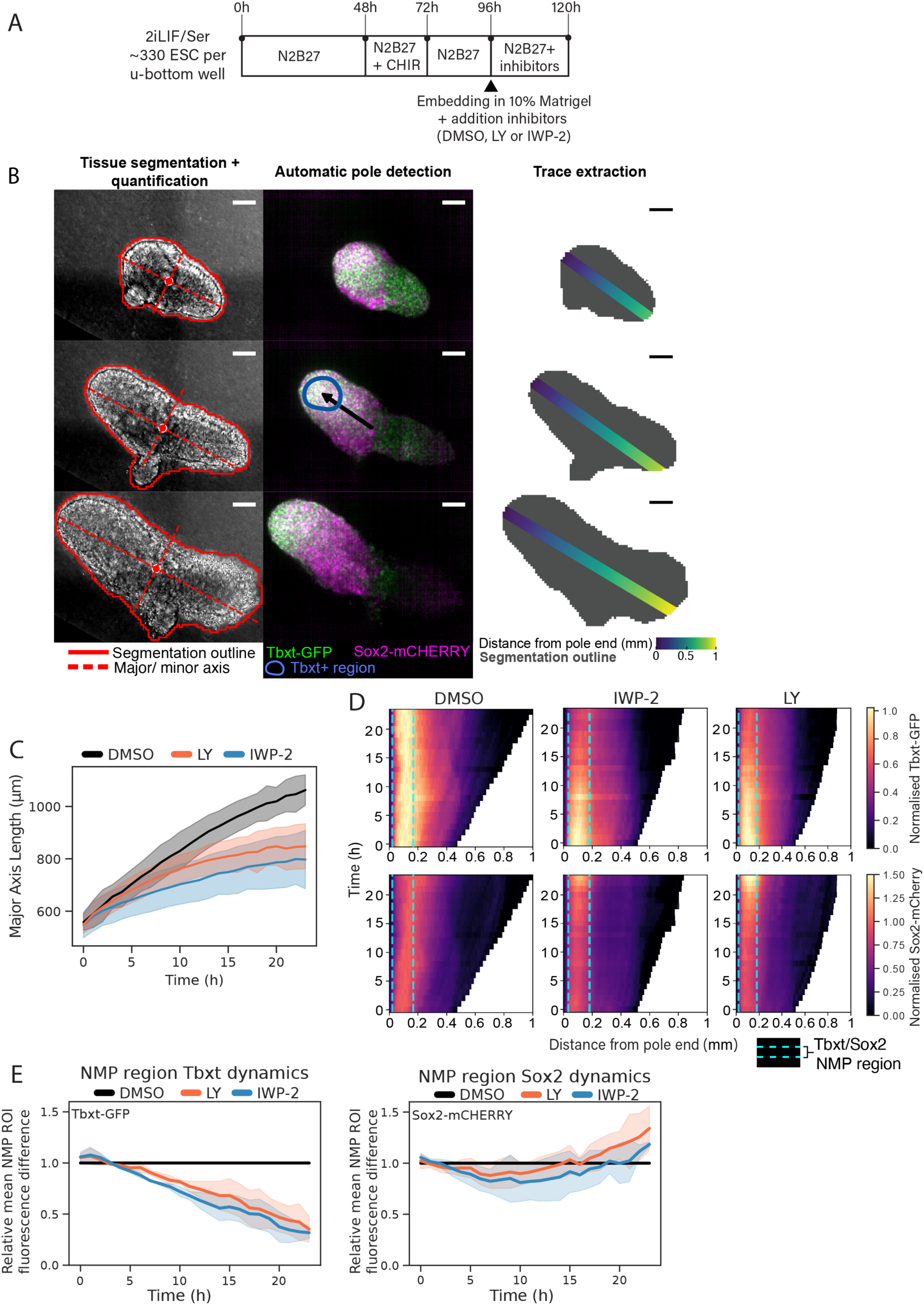
STR-KI cell analysis indicates that Tbxt-expressing cells are responsible for WNT and NOTCH driven STR-KI gastruloid elongation. **A.** Experimental setup for the generation of STR-KI gastruloids in 10% matrigel at 96 hours with DMSO, Wnt signalling inhibitor IWP2 (2.5 µM) or Notch signaling inhibitor LY411575 (150nM). Gastruloids were imaged on a confocal microscope every hour for 24 hours after embedding. **B.** Quantification pipeline of 3D gastruloid confocal image stacks. All scale bars indicate 100 µm. **C.** Axial length after treatment of gastruloids with IWP-2 or LY411575. **D.** Heatmaps showing the average Tbxt-GFP or Sox2-mCherry signal per binned distance from the pole along time post Matrigel embedding. **E.** Average difference in fluorescence signal of the Sox2/Tbxt region between control and treated conditions. N=3 independent experiments. Total gastruloids analysed: DMSO=35, LY= 38, IWP-2=41. Shaded areas indicate the mean +/- 0.95 confidence intervals, calculated by x̄ +/- z*(σ/sqrt(n)).

To determine how reporter expression responded to WNT or NOTCH inhibition during this period, whole gastruloid tissues were segmented and traces extracted along the extending long axis marked at the presumptive posterior end by Tbxt^GFP^ expression. Treatment with IWP2 or LY rapidly decreased Tbxt^GFP^ signal in the NMP region (within ∼2-3h). In contrast, Sox2^mCh^ signal in this region increased after ∼10-15 hours following an initial minor decrease compared to the DMSO control (Fig. 4D). This coincides in time with growth deceleration (Fig. 4D, E). Thus, the rapid Tbxt^GFP^ signal downregulation coupled with slowdown of axis elongation in response to WNT or NOTCH inhibition suggests AP axis elongation is driven predominantly by *Tbxt-*expressing cells.

### 5. Clonal tracing of in vitro derived NMCs reveals lineage bias of NMPs dependent on matrix and TBXT levels

Our findings at the bulk level suggest that differences in *Tbxt* levels influence lineage outcomes in Sox2/Tbxt co-expressing (NMC) subpopulations (Fig. 3D). However these experiments did not distinguish between neural or mesoderm lineage biases across the whole *in vitro* NMC-like population or a subset of already lineage-specified cells expressing the very highest or lowest reporter levels. To distinguish these possibilities, we sorted Sox2^mCh^pos;Tbxt^GFP^pos cells by FACS into low and high gates and deposited single cells into 96-well plates (Fig. 5A). We measured their differentiation outcome in FGF/CHIR, which supports both neural and mesodermal differentiation (Fig. 3D). A variety of substrates has been used in different NMC-generating protocols, and the influence of these is currently unclear. We therefore compared clonal outcomes on four commonly used matrices: fibronectin, gelatin, Geltrex and Matrigel. We then correlated this information with the lineage output using immunofluorescence and high throughput microscopy to quantify the SOX2/TBXT cellular phenotype in descendant cells in each well after five days (Fig. 5A, B).

**Figure 5.**
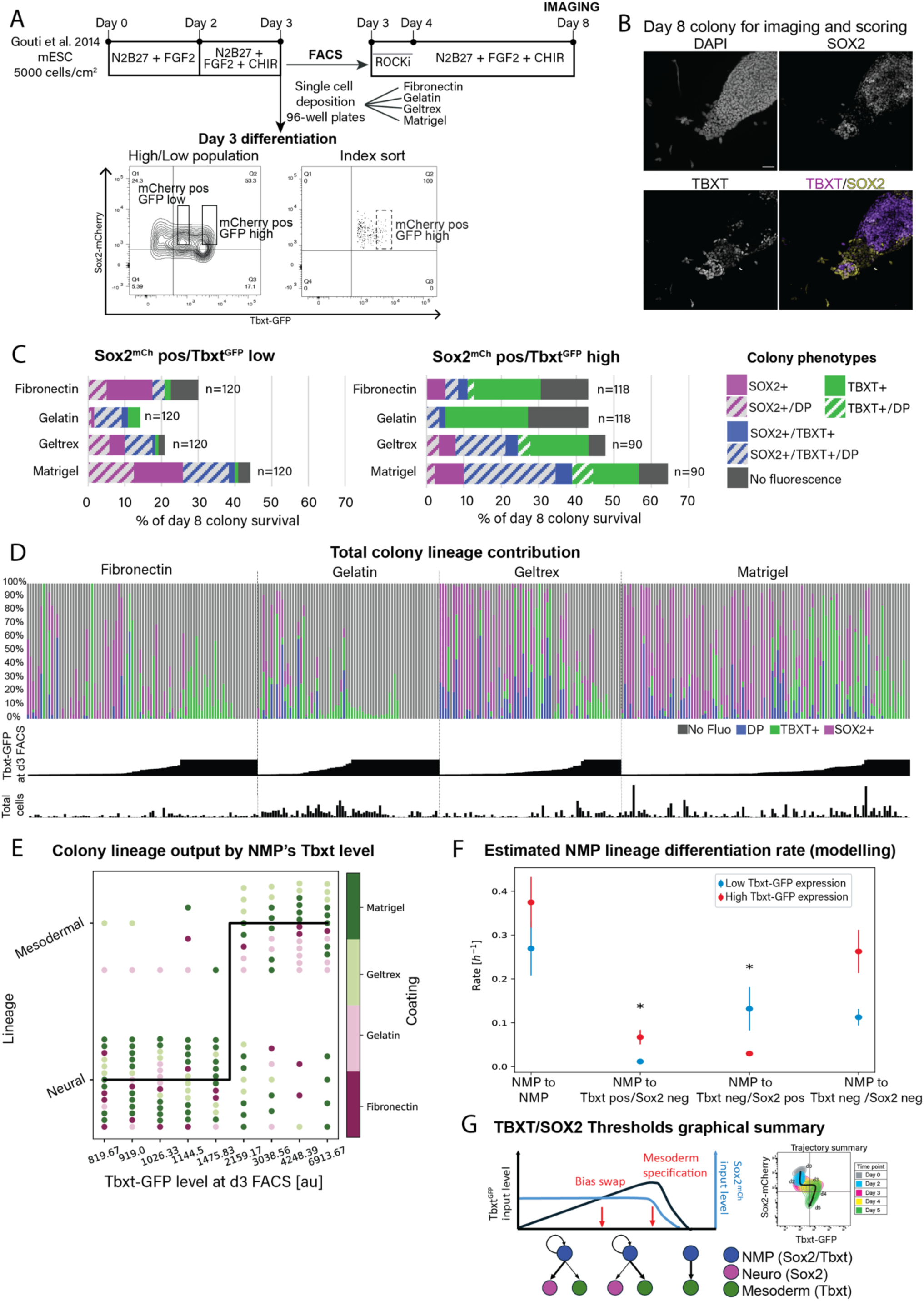
*In vitro* derived *Sox2*/*Tbxt* co-expressing cells are bipotent at clonal level and their lineage output is determined by *Tbxt* expression. A. Differentiation protocol scheme and representative gating strategy for FACS and index sorting according to Sox2^mCh^ and Tbxt^GFP^ expression. Cells were individually seeded into different substrates and cultured a further five days. Colony lineage output was assessed by immunofluorescence of endogenous SOX2 and TBXT proteins, after which the imaging data was segmented for individual cells in colonies and scored for their lineage output. **B.** Example of a colony on gelatin at the end of the experiment immunostained for SOX2 and TBXT. **C.** Total lineage output of colonies derived from single sorted Sox2^mCh^pos;Tbxt^GFP^high or Sox2^mCh^pos;Tbxt^GFP^low founder cells. Colonies were scored as containing SOX2pos, SOX2pos;TBXTpos (double positive (DP)), TBXTpos, and non-fluorescent (*i.e.* no signal detected) with n the number of colonies scored for each substrate. **D.** Overview of all individual colonies scored on their percent lineage output, classified by substrate and ordered by Tbxt^GFP^ level at the time of FACS (day 3). The total number of cells per colony ranged from 2-5335 cells. Note that the plateau in GFP level is arbitrary and corresponds to progenitors that were gated for Sox2^mCh^pos;Tbxt^GFP^high but not index sorted and demarcates a mesoderm commitment threshold. **E.** NMP lineage output as a function of initial Tbxt^GFP^ levels. Each dot represents one colony, coloured by substrate used: matrigel (dark green), geltrex (light green), gelatin (pink) and fibronectin (fuchsia). The Tbxt^GFP^ level on the X-axis is the middle value in the bin. **F.** Mathematical model to estimate fate in index sorted NMPs. The rate indicates whether single founder NMPs were above (red) or below (blue) the initial Tbxt level threshold of 1745 arbirtary units (au) as function of their colony output at the end of the experiment. The dots and the error bars correspond to the mean and the standard error of the mean, respectively. Asterisks indicate significant differences obtained after applying a Mann-Whitney statistical test (p < 0.05). **G.** ’two-threshold model’ summarising our findings.

We first considered the characteristics of all clones, sorted into ‘low’ and ‘high’ GFP gates as in Fig. 3A. In all clonal conditions tested, Sox2^mCh^pos;Tbxt^GFP^pos cells produced single positive neural or mesodermal descendants, as well as double positive cells, confirming that these cells are NMPs. Sox2^mCh^pos;Tbxt^GFP^low and Sox2^mCh^pos;Tbxt^GFP^high input cells showed distinct lineage output profiles. Sox2^mCh^pos;Tbxt^GFP^high cells gave rise to a greater proportion of Tbxt^GFP^ only colonies, confirming that high TBXT expression in individual Sox2^mCh^pos;Tbxt^GFP^high cells strongly predisposes them to a mesodermal outcome (Fig. 5C). From the population analysis (Fig. 3D), we can infer that double-negative descendants of Sox2^mCh^pos;Tbxt^GFP^pos cells are predominantly differentiated descendants of Tbxt^GFP^pos mesoderm. In these clonal assays, double negative cells were mainly produced by Sox2^mCh^pos;Tbxt^GFP^high cells, consistent with this interpretation. Interestingly, Sox2^mCh^pos;Tbxt^GFP^high cells showed higher plating efficiency than Sox2^mCh^pos;Tbxt^GFP^low cells, suggesting that increased TBXT promotes higher survival and/or adhesion to the substrate (Fig. 5C). Thus, at clonal level, high levels of GFP predict a high frequency of mesoderm-only outcomes, and an overall increased survival on multiple substrates.

Substrate composition also seemed to differentially affect cell phenotype. Strikingly, Matrigel and Geltrex reduced the overall mesoderm predisposition of Sox2^mCh^pos;Tbxt^GFP^high cells. This suggests that despite their propensity to differentiate towards mesoderm, Sox2^mCh^pos;Tbxt^GFP^high cells retain a substrate-dependent potential for neuromesodermal differentiation. Intriguingly, although the plating efficiency of both Sox2^mCh^pos;Tbxt^GFP^low and high cells was particularly poor on gelatin, this substrate appeared to enrich for growth of double positive cells from Sox2^mCh^pos;Tbxt^GFP^low cells (Fig. 5C, Suppl. Fig. 3C-E). Moreover, the substrate impacted the morphology of the colonies in culture: on gelatin, cells grew as compact colonies and produced an overall higher number of cells per colony, compared to Matrigel culture, where cells formed flat, dispersed colonies (Suppl. Fig. 3D). These observations suggest that once established from Sox2^mCh^pos;Tbxt^GFP^low cells, double positive cells are favoured by self-adherent growth. Lastly, we found that Sox2^mCh^pos;Tbxt^GFP^high colony survival on Geltrex and Matrigel was comparable, but Matrigel-coating increased the clone survival of Sox2^mCh^pos;Tbxt^GFP^low cells. Thus Sox2^mCh^pos;Tbxt^GFP^low and high cells exhibit differential behaviour that is modified by the substrate, which can alter both the cell phenotype and the extent of double-positive cell propagation.

We next compared individual colony composition with input reporter level (Fig. 5A, D). Cells in the Sox2^mCh^pos;Tbxt^GFP^high gate over two experiments predominantly produced only Tbxt^GFP^ positive mesoderm and/or double negative presumptive mesoderm (Fig. 5D, high GFP ‘plateau’ colonies). These two experimental repeats derived from populations where differentiation was relatively advanced and most cells lay in a Tbxt^GFP^ high;Sox2^mCh^ low state just prior to exit from the double positive population (Suppl. Fig. 3A, B – Exp 1-2). This suggests double positive cells reach a ‘mesoderm specification’ threshold beyond which mesoderm formation is virtually the only outcome and implying that some Tbxt^GFP^;Sox2^mCh^high cells are not NMPs, recalling our previous observation that, in vivo, a minority of TBXT/SOX2 immunofluorescent cells are present at the midline where we detect only mesoderm potential (Cambray & Wilson, 2007; French et al., 2024).

To investigate the relationship between input and output levels of Tbxt^GFP^ we then indexed a proportion of the input population, expanding the ‘Tbxt-high’ gate to include medium-level GFP-expressing cells, to assess whether the clone composition varies as a continuum or shows threshold responses to either reporter.

We used the indexed cells to mathematically determine whether the clonal lineage of individual NMP progeny is controlled by the initial levels of *Sox2* or *Tbxt* expression and the cell substrate. Sox2^mCh^ and Tbxt^GFP^ expression levels recorded at the time of plating (time = 0), were designated *S* and *T*, respectively. After five days, each seeded cell gave rise to varying numbers of cells (*n*) with four phenotypes based on their expression of TBXT and SOX2: *n_S-T+_*, *n_S+T-_*, *n_S+T+_* and *n_S-T-_*.

To determine whether the clonal lineage at day 5 post index sort was predominantly mesodermal or neural, we evaluated the difference between numbers of Sox2^mCh^ and Tbxt^GFP^ single positive cells per colony (*n_S-T+_*-*n_S+T_*_-_). The histogram of *n_S-T+_*- *n_S+T_*_-_ determined at time = 5 days, resembled a Gaussian distribution centred at *n_S-T+_*- *n_S+T_*_-_=zero (Suppl. Fig. 3F).

Of note, many NMPs had small *n_S-T+_*- *n_S+T_*_-_ values (close to zero) and it would be rather arbitrary to identify these NMPs as mesodermal or neural biased. Instead, the tails of the distribution should correspond with high confidence to mesodermal (*n_S-T+_* >> *n_S+T_*_-_) or neural (*n_S-T+_* << *n_S+T_*_-_) biased clones. Therefore, we used the standard error of the mean (SEM) of the distribution as a cut-off to assign NMP lineages, even though this excludes colonies from our analyses. Therefore, we classified the lineage of the progeny as mesodermal if *n_S-T+_* > *n_S+T_*_+_ by 27 cells (where 27 corresponds to the SEM). Conversely, we consider the clone to be neural biased when *n_S+T_*_-_ > *n_S+T_*_+_ by 27 cells. To investigate whether the lineage of the NMP progeny at time = 5 days depends on the Tbxt^GFP^ or Sox2^mCh^ level of the initial index sorted NMP cell at time = 0, we plotted the lineage classification of each seeded NMP as a function of both Sox2^mCh^ and Tbxt^GFP^ (Supp. Fig. 3G and Fig. 5E, respectively). To do this, we discretized the scales of Sox2^mCh^ and Tbxt^GFP^ by selecting bins of equal numbers of colonies.

The mesoderm or neural bias of the NMP progeny did not appear to be affected by the level of Sox2^mCh^ in the Sox2^mCh^pos;Tbxt^GFP^pos starting population (Suppl. Fig. 3G). In contrast, the starting level of Tbxt^GFP^ strongly influenced neural/mesodermal bias of the clones (Fig. 5D, E). At low values of Tbxt^GFP^, most NMPs gave rise to neural-biased clones (Fig. 5E). As Tbxt^GFP^ approached a value of 1745 arbitrary units in the index sort, the proportion of mesoderm-biased versus neural biased clones rapidly increased (Fig. 5D, E). Although the ratio of Sox2 to Tbxt has been cited as an important factor determining NMP fate (Koch et al., 2017; Morabito et al., 2025), this ratio did not predict the clonal outcome better than the levels of Tbxt^GFP^ alone (Suppl. Fig. 4 C-F). The substrate, however, did appear to modulate differentiation: neural-biased colonies were more abundant on fibronectin and Matrigel, whereas we did not observe a particular bias towards either lineage when cells were plated on gelatin or Geltrex of lineage-biased Sox2^mCh^pos;Tbxt^GFP^pos cells (Suppl. Fig. 3H). These results suggest the existence of a substrate-sensitive Tbxt^GFP^ expression threshold value (‘N-M bias swap threshold’) above which NMP cells are more likely to produce mesodermal cells, in addition to the mesodermal specification threshold described above.

To gain mechanistic insight into fate decisions in *in vitro* derived NMPs, we developed a minimal mathematical model of the proliferation and differentiation dynamics of individual NMPs (see Materials and Methods for details). We assumed that each NMP cell can divide symmetrically to: i) self-renew, giving rise to 2 NMPs (SOX2pos;TBXTpos); ii) differentiate into 2 mesoderm cells (SOX2neg;TBXTpos; iii) differentiate into neurectoderm (SOX2pos; TBXTneg); and iv) differentiate into double negative cells (SOX2neg;TBXTneg). Thus, our model has 4 parameters, which are the rates by which NMP cells give rise to each outcome. To check whether the model is sufficient to reproduce the dependence of the neural/mesodermal decision on the initial Tbxt^GFP^ levels in NMPs, we fitted our model to the experimental data.

When we grouped NMPs according to cell substrate, we observed that gelatin was associated with the highest rates of self-propagation of NMPs and their differentiation towards mesoderm (SOX2neg;TBXTpos), as well as descendants of nascent mesoderm, which are double negative (Suppl. Fig. 4A). Matrigel, on the other hand, produced the highest rate of differentiation towards neuroectoderm (SOX2pos; TBXTneg) (Suppl. Fig. 4A). Classifying NMPs according to whether they had Tbxt^GFP^ levels below or above the Tbxt^GFP^ threshold reported above (regardless of substrate), showed that Tbxt^GFP^ values above the threshold lead to a higher differentiation rate towards mesoderm, whereas Tbxt^GFP^ values below the threshold lead to neuroectoderm bias (Fig. 5F and Suppl. Fig. 4E), consistent with our results shown in Fig. 5E. Interestingly, for both populations, cells produced more NMPs at higher rates than either neural or mesoderm cells. This suggests that the threshold in differentiation bias does not affect the ability of NMPs to self-propagate and distinguishes this lower ‘N-M bias swap’ threshold from the high-Tbxt^GFP^ ‘specification threshold’, where cells predominantly exit the NMP state towards mesoderm differentiation. Despite the ability of fibronectin and Matrigel to bias cells towards neural differentiation, the mid-level threshold still holds when the experimental data are disaggregated according to the coating used (Suppl. Fig. 4B, F).

In summary, clonal analysis and mathematical modelling indicates that individual Sox2^mCh^pos;Tbxt^GFP^pos double positive cells *in vitro* are NMPs, and that threshold levels of *Tbxt* determine whether they are neural-biased or mesoderm-biased (N-M bias swap threshold). Levels of Tbxt^GFP^ in combination with Sox2^mCh^ determine mesoderm specification (mesoderm specification threshold; summarised in Fig. 5G). The cell substrate can modulate the proportions of neural versus mesoderm cells formed, while gelatin, a relatively challenging substrate for NMP adherence, once cells are attached, favours self-adherence and more efficient NMP self-propagation.

## Discussion

We generated a minimally disruptive embryonic stem cell line, STR-KI, that faithfully reports on *Tbxt* and *Sox2* expression. This cell line mirrors endogenous protein expression and minimises adverse phenotypic effects, lending confidence to the studies of lineage dynamics performed here. Reduction in SOX2 function negatively affects ESC maintenance and differentiation (Corsinotti et al., 2017), while reduction of *Tbxt* levels disrupts tail length (Gruneberg, 1958) and cell movement of *Tbxt* +/null cells in chimeras is defective (Wilson et al., 1995). Therefore, it is likely that normal protein function is preserved in this cell line since no abnormal phenotype or distribution of cells was detected *in vivo* or *in vitro*.

We further show that NMCs generated *in vitro* from mESCs (1) are bi-potent and (2) can self-renew in clonal analyses. These are functional attributes used to define *in vivo* NMPs (Tzouanacou et al., 2009). Our study of the differentiating population over time indicates that ESCs during NMP-like differentiation first reduce Sox2^mCh^ expression when exposed to FGF2, as expected in the epiblast *in vivo* (Corsinotti et al., 2017). Then, after CHIR addition, the double positive population (Sox2^mCh^pos;Tbxt^GFP^pos) emerges from these Sox2^mCh^ low cells, mimicking in vivo NMC emergence. Over the next 48 hours, Tbxt^GFP^ expression increases, until Sox2^mCh^ expression disappears from Tbxt^GFP^ high cells. Tbxt^GFP^ is later extinguished, resulting in a double-negative population. This cellular trajectory fits well with the known fate of epiblast cells that will eventually become NMPs *in vivo*: distal epiblast cells express relatively low *Sox2* levels and exit pluripotency to enter a bipotent NMP state in the caudal lateral epiblast or node-streak border. If NMPs encounter the primitive streak at the midline they will upregulate *Tbxt*, undergo epithelial to mesenchymal transition and exit the streak and its WNT-high environment as mesoderm, thereafter downregulating *Tbxt* (French et al., 2024; Kispert et al., 1995; Wymeersch et al., 2016).

*In vivo*, axial elongation is sensitive to perturbation of both WNT and NOTCH signalling, at least partly because NMPs require WNT for maintenance (Wymeersch et al., 2016) and both WNT and NOTCH for continued mesoderm production (Cooper et al., 2024; Martins-Costa et al., 2024; Wymeersch et al., 2016). TBXT is directly activated by WNT signalling (Yamaguchi et al., 1999), and optimal levels of TBXT require NOTCH signalling (Cooper et al., 2024). Our observation that inhibiting WNT or NOTCH signalling result in a decrease in TBXT is consistent with this published data. We show, however, that gastruloids undergo a particularly rapid slowing of axis elongation, concomitant with the suppression of Tbxt^GFP^ by either inhibitor (∼2-3h), while Sox2^mCh^ levels change only minimally for 10-15h. This suggests that adequate levels of both WNT and NOTCH are directly required to maintain TBXT-driven axial elongation.

We have shown that single *Tbxt/Sox2* co-expressing cells can differentiate towards SOX2+ neurectoderm and TBXT+ (and subsequently double-negative) mesoderm. This formally demonstrates that double positive cells generated using standard protocols from mESCs are indeed bipotent NMPs. Our mathematical modelling of index sorted founder cells and the fate in their respective colonies shows that higher Tbxt^GFP^ levels predispose cells towards mesoderm differentiation. Interestingly, this seems to occur at two distinct threshold levels of Tbxt^GFP^. Firstly, double positive cells with high Tbxt^GFP^ and diminishing Sox2^mCh^ fluorescence appear specified for mesoderm differentiation, reminiscent of a fraction of cells at the primitive streak which express high levels of TBXT and still express SOX2 but are not bipotent on transplantation to NM-fated regions (Cambray & Wilson, 2007; French et al., 2024). These cells are enriched in vivo for TBX6, a factor required to commit cells to a mesodermal fate (Takemoto et al., 2011). In support of this idea, TBX6 expression is elevated in TBXT^GFP^ high cells.

A second mid-level threshold of Tbxt^GFP^ appears to switch cells from a neural-biased to a mesoderm-biased state. This is somewhat unexpected because *in vivo* NMCs, even from predominantly mesoderm-fated regions (which express higher TBXT than those in neural-fated regions), can still produce neurectoderm on transplantation to more neural-fated regions (Wymeersch et al., 2016). However, our observation that cell substrate can modify the rate of differentiation suggests that the neural/mesodermal outcome of a clone/colony can also be influenced by environmental cues.

Unexpectedly, we did not find evidence that the ratio of Tbxt^GFP^ to Sox2^mCh^ affected the bias-swap threshold. This contrasts with previous studies suggesting that the Tbxt/Sox2 ratio or the ratio of WNT to Sox2 determines the outcome of differentiation (Koch et al., 2017; Morabito et al., 2025; Romanos et al., 2021). However, the levels of *Sox2* in the NMP compartment in those experiments are relatively constant. Indeed, in the mouse caudal lateral epiblast, the gradient of SOX2 expression is much shallower compared to that of TBXT (French et al., 2025). If the TBXT:SOX2 ratio determined NMP outcome, loss of TBXT function would be expected to bias cells towards neural differentiation. However, *in vivo*, chimaera analysis of TBXT^-/-^ cells does not seem to show this bias (Guibentif et al., 2021). Therefore, it is conceivable that against a baseline of relatively uniform SOX2, the TBXT:SOX2 ratio might appear to predict neural versus mesodermal outcome, when TBXT levels might be the main determinant of differentiation bias. Nevertheless, SOX2 may serve an important function in the NMC population. In zebrafish, ectopic expression of SOX2 in NMCs prevents their exit to the mesoderm (Kinney et al., 2020). Together with the observation that TBX6 activation upon entry to the primitive streak suppresses SOX2 to prevent ectopic neural differentiation in the paraxial mesoderm (Takemoto et al., 2011), this suggests SOX2 threshold levels operate at the point of exit of cells from the NMC population. Interestingly, ordering *in vivo* single NMP transcriptomes along a differentiation trajectory shows that peak *Tbxt* mRNA levels occur in double-positive cells, and *Tbx6* mRNA, signifying mesodermal commitment, begins to accumulate only in cells with lower levels of both mRNAs (Gouti et al., 2017). Our observation that during in vitro NMP differentiation, TBXT^GFP^ increases while Sox2^mCh^ levels remain constant, then the levels of both reporters drop before cells exit the double positive population is consistent with such a gatekeeper function of SOX2, but not with a continuously variable mutual inhibition of SOX2 and TBXT.

The primacy of TBXT dosage in influencing the outcome of differentiation recalls previous experiments *in vivo* where levels of TBXT dictate cell behaviours: low-TBXT expressing cells stay in the primitive streak/tail bud,and exit as mesoderm if TBXT levels are increased (Wilson & Beddington, 1997; Wilson et al., 1995). Our study indicates that this works at the individual cell level, whereby cells sense cues in their environment (e.g. WNT, NOTCH), respond by tuning TBXT up or down, which in turn influences the differentiation outcome at the single cell level. Furthermore, quantitative reduction in wildtype TBXT gene dosage progressively shortens the axis (Gruneberg, 1958; Stott et al., 1993; Xia et al., 2024). This suggests that continued axial elongation (through the persistence of NMPs) is sensitive to TBXT levels through modulating these probabilistic states. Since gastruloid manipulation during elongation shows this TBXT sensitivity, such stem-cell based models will prove useful in the future to investigate the mechanistic bases of these molecular thresholds during developmental processes, such as axis elongation.

## Materials and methods

### Reporter mESC generation

To generate the double reporter cell line, we performed CRISPR-based transgenesis in two targeting rounds (one for each gene locus, starting by *Tbxt*) in E14tg2a ESCs following an established CRISPR-Cas9 protocol (Dewari et al., 2018). crRNA and sgRNA guides are listed in Table S1. Briefly, we replaced the stop codon at the C-terminus of the targeted genes by homologous recombination with a reporter construct containing T2A-mCherry (in the case of *Sox2*) or P2A-H2B-GFP (in the case of *Tbxt*) (Fig.1A). The *Tbxt* targeting vector contained a puromycin resistance cassette, which was used for clone selection. *Sox2*+ cells were selected by FACS, sorting individual mCherry+ cells into 96-well plates. Single clones were picked, expanded, and verified for correct transgene integration by PCR and nanopore sequencing (Suppl. Fig. 1A, B).

### PCR and nanopore sequencing

Correct integration of the fluorescent protein sequences into the endogenous loci of *Tbxt* and *Sox2* was confirmed by PCR amplification of the targeted site. The primer sequences used are listed in Table S1. To validate the sequence, PCR products were subcloned into a plasmid using the Zero Blunt™ TOPO™ PCR Cloning Kit and sequenced via Nanopore sequencing.

### Mouse husbandry

C57BL/6 female mice (Charles River) were used for chimaera generation. MF1 mice were used for electroporation experiments. ICR wildtype mice (JAX stock #009122, The Jackson Laboratory) were used for immunostaining. Mice were maintained on a 12 hr-light/12 hr-dark cycle. To collect embryos at specific developmental stages, timed matings were set up overnight. Noon on the day of finding a vaginal plug was designated as embryonic day (E) 0.5. Embryos dissected for immunostaining were approved by the Ethical Committee of the RIKEN Center for Biosystems Dynamics Research (A2016-03-13).

### Chimaera generation

C57BL/6 female mice were superovulated (100 IU/mL PMSG and 100 IU/mL hCG intraperitoneal injections 48 h apart) and crossed with wild type studs. Pregnant mice were culled at E2.5 by cervical dislocation, ovaries with oviducts were dissected and collected in pre-warmed M2 medium. Oviducts were flushed using PBS and a 20-gauge needle attached to a 1 mL syringe and filled with PB1 (Whittingham, 1974). E2.5 embryos were collected and washed in PB1, their zona pellucida removed using acidic Tyrode’s solution and transferred to a plate with incisions where of 8-15 cells were added to each embryo. Embryos were then incubated at 37°C in 5% CO2 for 24h prior to transfer to pseudopregnant females. Blastocysts were selected and collected to be transferred into the uterus of a pseudopregnant CD-1 female. Females were culled according to Schedule 1 (Animals (Scientific Procedures) Act 1986), and embryos were dissected at E6.5-9.0 in M2 medium and observed for chimeric ESC contribution under an Olympus IX51 microscope. The sex of embryos used in this study was not determined. All reagents are listed in Table S1.

### Focal electroporation and fate mapping

A more detailed description can be found in (Huang et al., 2015). E7.5 neural plate (NP) stage embryos were dissected in M2 medium with their extraembryonic cavities intact, but Reichert’s membrane removed. A PBS-filled 30 mm petri dish was prepared under a stereomicroscope (Zeiss Stemi 2000-C) with a PBS-filled, capillary-sheathed, platinum point electrode (anode; final diameter of capillary sheath 20-30 µm) and an 0.2mm L-shaped platinum wire (cathode) connected to an ECM 830 square wave pulse generator (BTX). Using a pneumatic pico pump (World Precision Instruments, PV830) and micropulled injection needles, a small volume (<5µL) of DNA solution (pCAG::GFP plasmid at 1-1.5 μg/mL, 0.01% Fast Green food colouring dye in PBS) was injected in the amniotic cavity of an NP-stage embryo. Embryos were briefly transferred from M2 medium to the PBS-filled electroporation dish, and electroporated using 200 Volts in 6 pulses, each of 50ms duration with a 1s interval between each pulse. Embryos were then immediately transferred to pre-equilibrated culture medium and cultured in 4-well plates in an incubator supplied with 5% CO2 in air at 37° C for 24h (Cambray & Wilson, 2007). GFP contribution was assessed at 2h post-electroporation using a fluorescence compound dissecting microscope (Nikon AZ100). For assessment of contribution, samples were fixed, immunostained as wholemount as described below using an anti-GFP primary antibody (Abcam ab13970, 1:800) and Liquid DAB+ Substrate Chromogen System (Dako K3467). Imaging and scoring of contribution were done under a Nikon AZ100 microscope.

### Immunofluorescence on wholemount embryos

E6.5-E8.0 embryos were dissected in home-made M2 medium (Nowotschin et al., 2010), their yolk sac and amnion membranes removed, after which they were fixed for 25min at 4°C in 4% PFA solution (Nacalai Tesque #09154-85). After three short washes in PBS in 0.1% Triton X-100 (now referred to as PBST), samples were permeabilized in 0.5% Triton X-100 in PBS for 15 min, followed by 20min in 0.5 M glycine in PBST, two 10min washes in PBST and blocked overnight at 4°C in 10% donkey serum (Merck) in PBS/0.3% TritonX100. Primary antibodies were diluted in blocking buffer for 48h on a rocking platform at 4°C. Antibodies used (supplier, final concentration): anti-Sox2 (Abcam, ab92494, 1:200); anti-TBXT (R&D, AF2085, 1:200). After two days, four 25min washes were performed with PBST on a rocking platform at room temperature. Alexa Fluor®-conjugated secondary antibodies were diluted in blocking buffer at 2 µg/ml final concentration (Thermo Fisher Scientific) and incubated for 48h at 4°C, then washed in PBST (4x25min) with the last wash containing DAPI (Invitrogen, D3571, final concentration at 2.5 µg/mL). To image, samples were cleared in BA:BB (2:1 benzyl alcohol:benzyl benzoate; Sigma) after dehydration through an increasing methanol (Nacalai Tesque) /PBS steps (25%, 50%, 75%, 2x 100%, 5min each).

Wholemount embryo samples were imaged under a LSM800 confocal system using GaAsP detectors (Zeiss). Three replicate embryos were imaged per developmental stage. Whole-mount immunostaining data was processed using Zeiss software (Zeiss). For DAPI channel processing, a median filter and background subtraction were applied. Orthogonal projections were generated as X-Y weighted averages of 8 consecutive slices, representing a total of 8 µm across the Z-axis. IMARIS software (Oxford Instruments) was used to generate 3D renderings.

### Cell culture, differentiation and flow cytometry/FACS

STR-KI mESCs were maintained on 0.1% gelatin-coated plates in Glasgow Minimum Essential Medium (GMEM, Sigma-Aldrich #G5154) supplemented with 10% foetal calf serum (Gibco #10270-106), 100 U/mL LIF (made in-house), 100 µM 2-mercaptoethanol (Gibco #31350-010), 1X non-essential amino acids (Gibco #11140-035), 2 mM L-Glutamine (Invitrogen #25030-024) and 1 mM Sodium Pyruvate (Invitrogen #11360-039). mESCs were passaged every other day using 1X trypsin (Sigma #59429C) after washing with PBS.

For NMP differentiation, following Gouti et al. 2014, STR-KI mESCs were plated at a density of 5,000 cells/cm^2^ on 0.1% gelatin-coated plates in N2B28 medium. N2B27 medium consisted of a 1:1 mixture of Neurobasal (Thermofisher #21103049) and Advanced DMEM/F12 (Thermofisher #12634028) supplemented with 2 mM L-Glutamine, 50 µM 2-mercaptoethanol, 0.5X B27 (Thermofisher #17504001) and 0.5X N2 (Thermofisher #17502048). Cells were supplemented with 10 ng/mL FGF2 (R&D Systems #3718-FB) for 48 hours, and for 24 additional hours with both 10 ng/mL FGF2 and 5µM CHIRON99021 (Axon #1386).

For flow cytometry analyses, after TrypLE Express Enzyme 1X treatment and centrifugation, cells were resuspended in PBS containing 2% FCS and 0.1 µg/mL Draq7. For FACS resuspension was done in N2B27 medium containing 0.1 µg/mL Draq7.

Cells were analysed using a 4 laser LSR Fortessa (BD) flow cytometer, employing B 530/30-A, Y/G 610/20-A, and R780/60-A laser/filter combinations. For cell sorting, a BD FACS Aria II Cell Sorter was used with B 525/50-A, Y/G 610/20-A, and R780/60-A laser/filter combinations.

For single cell plating experiments, differentiated cells were detached using TrypLE Express Enzyme 1X (Gibco 12604013) and sorted into CellCarrier 96-well plates coated with one of the following substrates: 0.1% gelatin, fibronectin (16.6 µL/mL in PBS; Sigma, #F1141), Matrigel (200 µg/mL in PBS; Corning, #354277), or Geltrex (120 µg/mL in Advanced DMEM/F12; Thermofisher, #A1413201). Plates were coated and incubated for 1 hour at 37°C prior to cell seeding. Cells were cultured in N2B27 supplemented with 100 U/mL penicillin/streptomycin (Invitrogen, #15140-122), 10 µM ROCK inhibitor Y-27632 (Tocris, #1254/10) and either 10 ng/mL FGF2 or in combination with 5µM CHIRON99021. After 24 hours, the medium was replaced without ROCK inhibitor. The ROCK inhibitor Y-27632 was used to enhance single-cell survival (Watanabe et al., 2007) and was used in FACS experiments to counteract dissociation and cell-stress-induced apoptosis.

All cells were maintained under standard culture condition (37° C and 5% CO2).

### Quantitative reverse transcription polymerase chain reaction (qRT-PCR)

A total of 200 cells were sorted into PCR tubes containing 10 µL of 2X Reaction Mix from the CellsDirect One-Step qRT-PCR Kit (Invitrogen #11753100) with 0.4 U/µl of RNAse inhibitor (Invitrogen #AM2694) and flash-frozen on dry ice. Reverse transcription reactions were performed using the SuperScript III RT/Platinum Taq mix provided in the same kit. qPCR was performed using LightCycler 480 SYBR Green I Master (Roche, #4887352001) in 384-well LightCycler 480 Multiwell Plates (Roche #04729749001) and using the Roche LightCycler 480 Real-Time PCR System. The *Tbp* gene was used as an endogenous control, and relative gene expression was calculated using the 2^-ΔΔCt^ method.

### Immunocytochemistry and confocal microscopy

Cells were fixed with 4% paraformaldehyde for 10 minutes, then washed twice with PBS over a total of 10 minutes and permeabilised with 0.5% Triton X-100 in PBS for 10 minutes. Following two additional PBS washes, cells were blocked for 30 minutes in a solution containing 5% donkey serum and 0.1% Triton X-100 in PBS. Primary antibodies, diluted in the same blocking solution, were added overnight at 5°C. After washing twice with PBS, secondary antibodies, diluted in blocking solution, were added for 2 hours at room temperature. After two further PBS washes, DAPI (Invitrogen, D3571, final concentration at 2.5 µg/mL) was added for 5 minutes at room temperature, followed by two PBS washes. Imaging of multi-well plates was performed using the High-Content Imaging Opera Phenix Plus from the IRR Imaging Core Facility. Single cell colony lineage scoring was performed using Signals Image Artist software (Revvity), using the 3D Analysis stack processing pipeline. Image stacks were processed to identify and segment colonies. For cell counting, the Find Nuclei – Method C function was utilized. Following segmentation, both cell number and fluorescence intensity data were extracted from the analysed image sets.

### Gastruloid culture, live imaging and quantitative analysis

Gastruloid culture was carried out as previously described (van den Brink et al., 2020 Vianello et al., 2020 [dx.doi.org/10.17504/protocols.io.9j5h4q6]). In summary, 2i/LIF/FCS mESCs were washed in PBS before adding 0.05% Trypsin EDTA solution. After cell detachment, 5-10 volumes of fresh 2i/LIF/FCS were added to quench the Trypsin EDTA solution. Cells were transferred to a universal tube and pelleted by centrifugation at 300g for 3 minutes. The media was aspirated, and the pellet was resuspended in cold PBS to make a single cell suspension. A PBS wash was repeated once more, then the cell pellet was resuspended in prewarmed N2B27 to a single cell solution. Cells were counted and diluted to a solution of 8,250 cells/mL in N2B27 medium, after which 40μL of this medium (containing ∼330 cells) was plated into untreated u-bottom 96 wells and incubated for 48 hours at 37°C and 5% CO2. At 48 hours, an additional 150μL of N2B27 and 3 µM CHIR99021 were added to the well. The last half of the additional media was expelled forcefully to dislodge the aggregate but without spilling the medium. The aggregates were cultured for a further 24 hours to d3, after which 150μL of media was removed and replaced with 150μL fresh N2B27. At d4, gastruloids were washed in N2B27 and then transferred to flat-bottom and black-walled 96-well plates (PerkinElmer) containing ice-cold phenol-red free N2B27 (Thermofisher #21041025, #12348017) + 10% matrigel supplemented with LY 411575 (150nM), IWP-2 (2.5μM), or an equal amount of DMSO. Gastruloids were left for 5 minutes to drop to the bottom of the well and then placed into an incubation chamber at 37°C and 5% CO2 to set the matrigel. After this period, gastruloids were imaged on a confocal microscope (Opera Phenix Plus) every hour for 24 hours. Transmitted light and fluorescence signal was collected using epifluorescence and a 20x objective. Excitations of 488nm and 561nm, emissions of 522nm and 599nm, and exposure times of 0.1s and 0.2s were used for GFP and mCherry respectively. XYZ voxel dimensions were 0.89x0.89x5 microns.

Image analysis was performed in python using custom scripts and open-source libraries that can be found at 10.5281/zenodo.17192095. Whole tissue segmentation masks were generated using with the Segment Anything Model (SAM) (arXiv:2304.02643). Extending poles of gastruloids were inferred by median projecting Tbxt^GFP^ signal z-stacks to 2D, smoothing with a Gaussian filter, and isolating the largest domain of pixels over the 99th percentile. The vector between the centroid of this domain and the centroid of the whole gastruloid mask determines the orientation to the *Sox2/Tbxt* co-expressing region. The orientation calculated at 10 hours post embedding was applied to all other timepoints. Traces were collected in a 90μm wide linear line orthogonal to the vector orientation. Extending pole “ends” containing Sox2/Tbxt coexpressing domains were isolated as a bin between 35μm and 178μm from the end of the pole. The limits of this region were manually determined by the overlap of Tbxt^GFP^/Sox2^mCh^ signal gradients and previous observations of the location of NMP regions in vivo (i.e the tail tip) (French et al., 2025). Signal of Tbxt^GFP^ and Sox2^mCh^ was measured in 2D by the average intensity from median z-projections of both channels in 0.89μm wide bins along the gastruloid trace. Channel signal was normalised per gastruloid to the 5^th^ and 99^th^ percentile of trace signal values at 0h, then per timepoint and condition normalised to the 99^th^ percentile of DMSO control. For the NMP region Tbxt/Sox2 dynamics, the difference between signal from treated and control conditions was calculated per replicate and timepoint.

For flow cytometry analyses, an average of 60 gastruloids were pooled per experiment and bulk analysed at 72hpa and 30 gastruloids at 96hpa. Analyses were done using the NovoCyte Penteon analyser (Agilent). Flow data was analysed in R using the CytoExploreR plugin (Dillon Hammill (2021), CytoExploreR: Interactive Analysis of Cytometry Data. R package version 1.1.0. https://github.com/DillonHammill/CytoExploreR).

For imaging, gastruloids were fixed with 4% PFA for 40 min at room temperature and stained with DAPI overnight. Gastruloids were mounted in low melting point agarose in a 96-well ibidi plate and clarified with RapiClear 1.49 overnight. An average of 3-5 gastruloids per condition were imaged using an OperaPhenix confocal microscope, a 300 µm section was imaged in 2 µm steps and representative examples were selected. Image analysis was performed in Fiji. E14tg2a gastruloids were used as a negative control to set fluorescence thresholds. A MAX intensity projection of the DAPI signal was used to create outlines of the gastruloids using a custom Fiji macro.

### Mathematical modelling

In this study, we propose a minimal mathematical model of the dynamics of NMP cells and their progeny to investigate the cell fate decision process. We assume that each NMP cell divides symmetrically, resulting in i) two NMPs, or ii) two cells that are simultaneously Tbxt+ and Sox2-, or iii) two cells that are simultaneously Tbxt- and Sox2+, or iv) two cells that are simultaneously Tbxt- and Sox2-. We also assume that, with the exception of NMPs, the other cells do not divide within the time window of our study. The model is encoded in the following linear system of ordinary differential equations describing the time course of NMPs and their progeny:

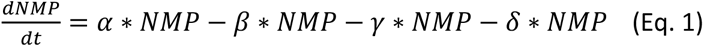

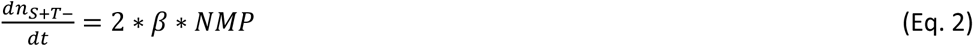

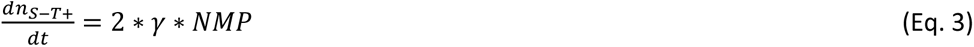

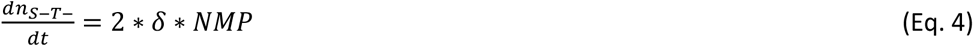

Where *t* is time, *NMP*, 𝑛*_S+T_*_–_, 𝑛*_S–T_*_+_and 𝑛*_S–T_*_–_ are the number of NMPs, the number of cells that are simultaneously Sox2+ and T-, the number of cells that are simultaneously Sox2- and T+ and the number of cells that are simultaneously Sox2- and T-, respectively. The model has 4 parameters: 𝛼, 𝛽, 𝛾 and 𝛿, which correspond to the rates at which NMP cells give rise to more NMP cells, cells positive for Sox2 but negative for T (*i.e*., neural lineage), cells negative for Sox2 but positive for T (*i.e.*, mesodermal lineage) and cells negative for both Sox2 and T (double negative).

To test whether the model was sufficient to reproduce the dependence of the neural/mesodermal decision on the *T* level of the NMP, we fitted the model to the experimental data.

This system has an analytical solution given by

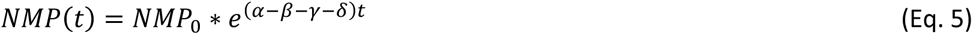

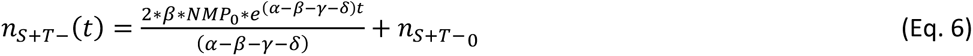

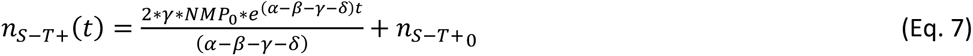

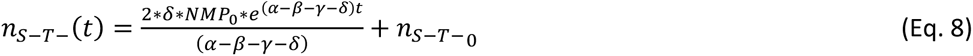

Where *NMP_0_* is the initial number of NMP cells, 𝑛*_S+T–_*_0_ is the initial number of Sox2 positive but T negative cells, 𝑛*_S–T+_*_0 i_s the initial number of Sox2 negative but T positive cells and 𝑛*_S–T+_*_0_ is the initial number of double negative cells.

Finally, we assume the following initial condition, reflecting the experiments described in section 3.

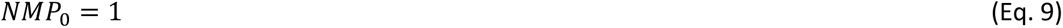

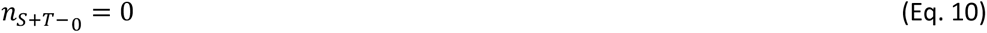

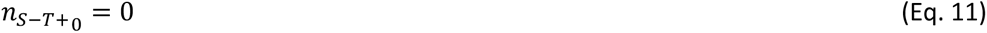

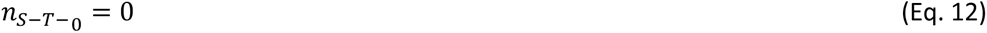

To fit the model to the experimental data, we minimized the distance function δ between the experimental number of cells and the model predicted number of all cell types for each experiment. The distance d was defined as:

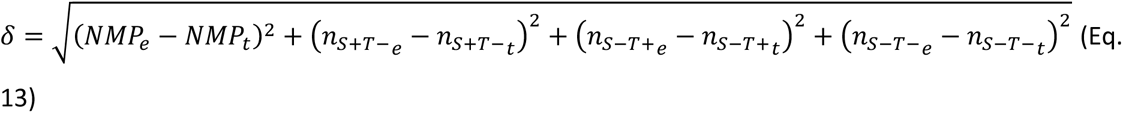

Where *NMP_e_* and *NMP_t_* are the experimental and model predicted number of NMP cells, 𝑛*_S+T–_e__* and 𝑛*_S+T–_t__* are the experimental and model predicted number of cells positive for SOX2 but negative for TBXT, 𝑛*_S–T+_e__* and 𝑛*_S–T+_t__* are the experimental and model predicted number of cells positive for T but negative for SOX2, and finally 𝑛*_S–T–_e__* and 𝑛*_S–T–_t__* are the experimental and model predicted number of cells negative for both SOX2 and TBXT. From all best-fitting parameter values that minimized d for each experiment, we calculated the mean and standard error of the mean.

A Jupyter Notebook (http://jupyter.org/) containing the source code used to generate the mathematical model plots in Fig. 5 and Suppl. Fig. 3, 4 can be found at 10.5281/zenodo.17154933 (Ceccarelli & Chara, 2025).

## Author contributions

Conceptualization: A.B-C., V.W;

Methodology: A.B.-C., A.G, A.C, O.C, V.W;

Formal analysis: A.G, A.C, O.C, M.F

Investigation: A.B.-C., A.G, A.C, E.K, F.J.W, R.P, M.F, J.A, Y.H, A.S.B, D.L.R,

Resources: F.C.K.W, F.J.W, M.T, S.L

Data curation: A.G, A.B-C, V.W. O.C, A.C;

Writing - original draft: A.B-C, V.W;

Writing - editing: A.B-C, V.W, F.J.W, S.L, M.Y, A.G, F.C.K.W, A.C, O.C;

Visualization: A.B.-C., A.G, A.C, F.W, M.F, J.A, V.W

Supervision: A.B-C, O.C, V.W;

Project administration: A.B-C, V.W;

Funding acquisition: A.B-C, V.W

## Acknowledgements

The authors thank the IRR core facilities at the University of Edinburgh for their important contributions to this work, particularly the IRR flow cytometry, tissue culture and high content imaging facilities. We thank Augusto Borges for technical advice on modelling aspects of the study. We would also like to thank André Dias for critically reading an early version of the manuscript.

## Data availability

Data availability: All relevant data can be found within the article and its supplementary information.

---

**Supp. Figure 1.**
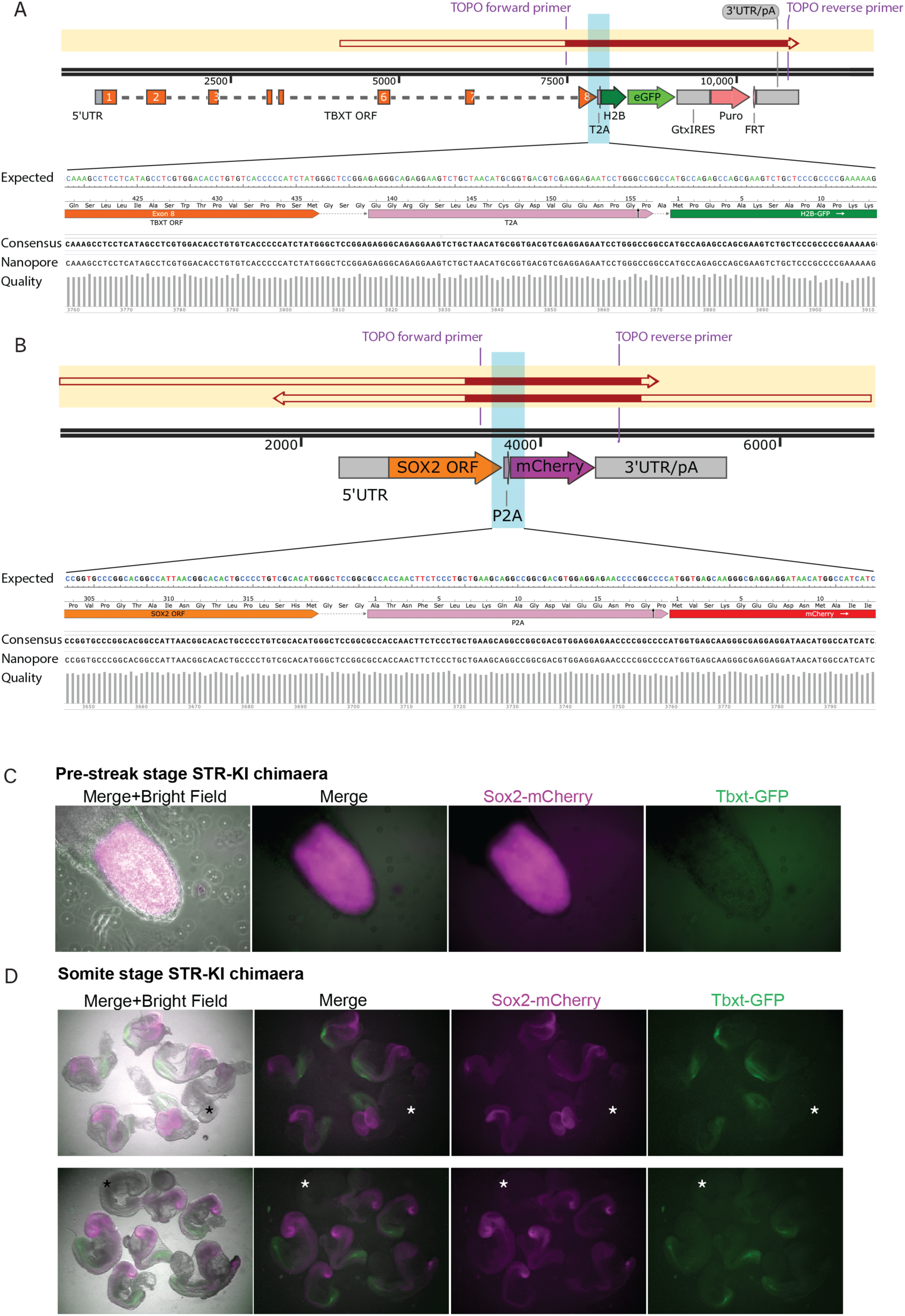
STR-KI cell line contributes highly to chimaeras. A-B. Schematics of targeted region of the Tbxt **(A)** and Sox2 **(B)** loci in the STR-KI cell line showing correct integration and sequence alignment to the consensus after nanopore sequencing. **C.** mouse pre-streak and **D.** E8.5 (2-12) somite stage chimeras appear morphologically normal, show high contribution of STR-KI cells, and correct location of the reporter fluorescent proteins. Asterisk in D shows a non-contributing littermate as a negative control.

**Supp. Figure 2.**
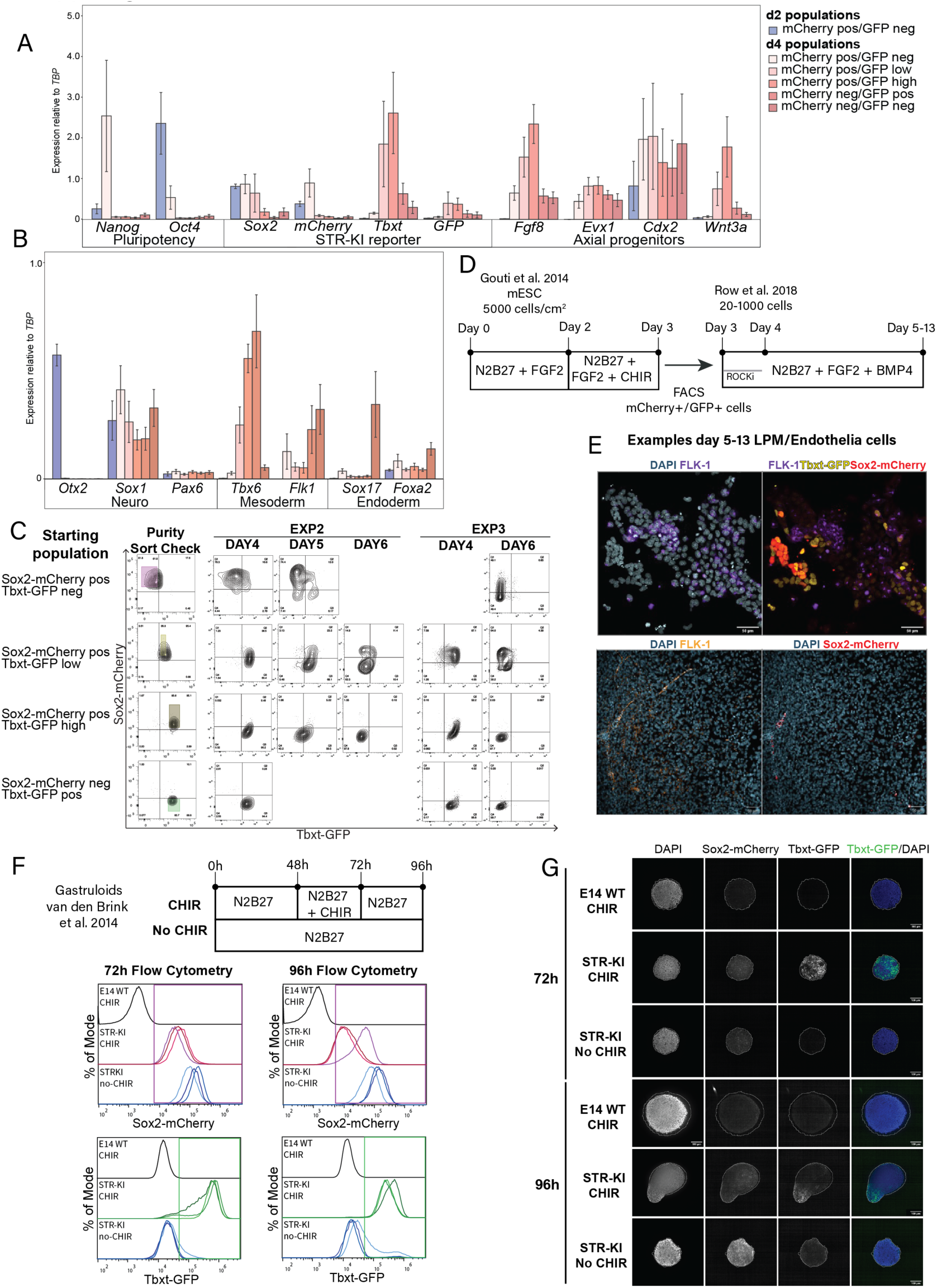
*In vitro* derived *Sox2*-mCherry and *Tbxt*-GFP co-expressing cells are NMPs, and their lineage trajectory can be captured for potency studies. A-B. qRT-PCR analysis of populations on day 2 and day 4 using the same gating strategy and differentiation protocol in Fig. 2A. Gene markers of pluripotency, axial progenitors (inset A), and neural, mesodermal, endodermal differentiation markers (inset B) with n=3 independent experiments for each population. **C.** Repeats of the time-course experiment described in Fig. 2C-D. A factor of variability between these three experiments can be noted and is inherent to any differentiation protocol. However, cell populations of all three experiments followed the same trajectory trends as summarised in Fig. 2D. A representative purity check of the initial FACS populations is shown. **D.** Scheme of the experimental set up for FACS *Sox2*^mCherry^*Tbxt*^GFP^ double positive progenitors and testing of lateral plate mesoderm (LPM) potency. **E.** Colonies stained for the LPM marker Flk-1. In some Flk-1+ colonies changes in morphology where minimal (top) whereas in others elongated FLK-1+ cells were observed, indicating potentially more progressed endothelial differentiation. **F.** Experimental setup for the generation of gastruloids to test symmetry breaking and *Tbxt*^GFP^ activation upon CHIR activation. Flow cytometry on an average of 60 gastruloids were pooled and bulk analysed for 72 hpa and 30 gastruloids for 96 hpa (n=1). **G.** Immunostaining of wildtype (WT) or STR-KI derived gastruloids for *Sox2*^mCherry^ and *Tbxt*^GFP^ at 72 and 96 hpa. Note the absence of Tbxt^GFP^ in both WT and no CHIR treated gastruloids. Axis elongation is visible between 96-120 hpa, depending on the cell line use, with our STR-KI reporter showing elongation from 96 hpa upon CHIR treatment representative image of 3-5 gastruloids analysed per sample.

**Supp. Figure 3.**
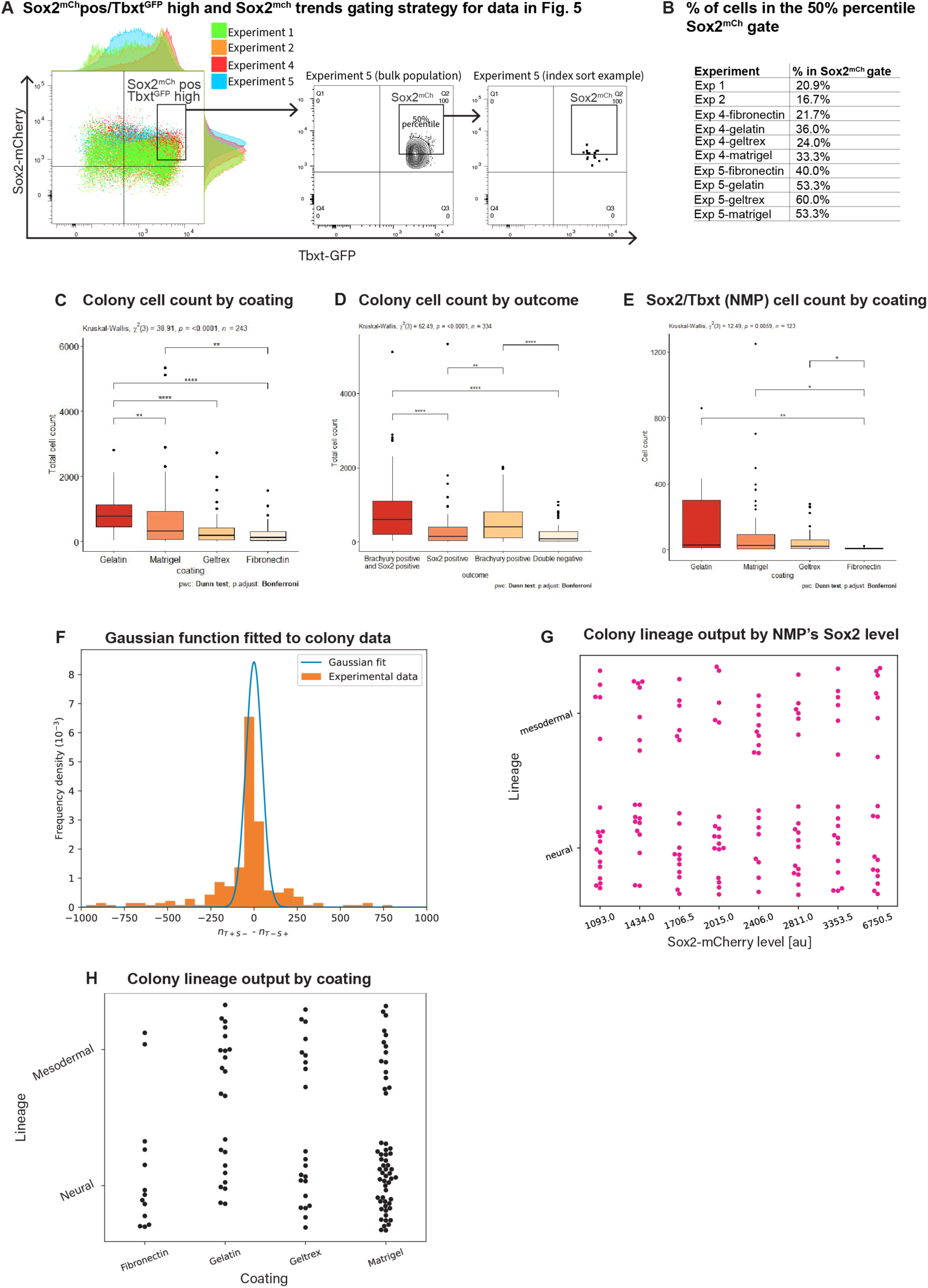
Quantitative analysis and mathematical modelling of clonal data. A. Overlap of bulk-flow data from NMP differentiations where single cells that fell in the Sox2mChpos/TbxtGFPhigh sorting gate were reanalysed by level of Sox2. Exp 1,2 were unindexed, and Exp 4,5 were indexed. The % of cells above the 50% percentile of mCherry were calculated based on the least advanced differentation repeat (exp5). **B.** Summary table listing the % of cells within the Sox2mCh/TbxtGFP high population falling in the top 50% percentile of Sox2mCherry across all experiments. **C.** Total number of cells per colony measured after clonal expansion of single cells grown on different extracellular matrices or classified by colony phenotype (**D**). **E.** Total number of SOX2+TBXT+ cells per colony cultured on different coating matrices. **F.** Histogram for *n_S-T_*_+_ - *n_S+T_*_-_ (orange bars). Blue line: Gaussian function fitted to the data. **G.** Clonally plated Sox2 ^mCherry^ Tbxt^GFP^ (NMP) lineage output as a function of Sox2^mCherry^ levels in individually plated NMPs. Each dot represents one colony. The Sox2^mCherry^ level on the X-axis is the middle value in the bin. **H.** Clonally plated Sox2 ^mCherry^ Tbxt^GFP^ (NMP) lineage output as a function of the extracellular coating used. For C-E: Dots are the mean with error bars, the standard error of the mean. Asterisks indicate statistically significant differences (*p* < 0.05) obtained after applying a Mann-Whitney statistical test (* p < 0.05, ** p < 0.01, *** p < 0.001, **** < 0.0001).

**Supp. Figure 4.**
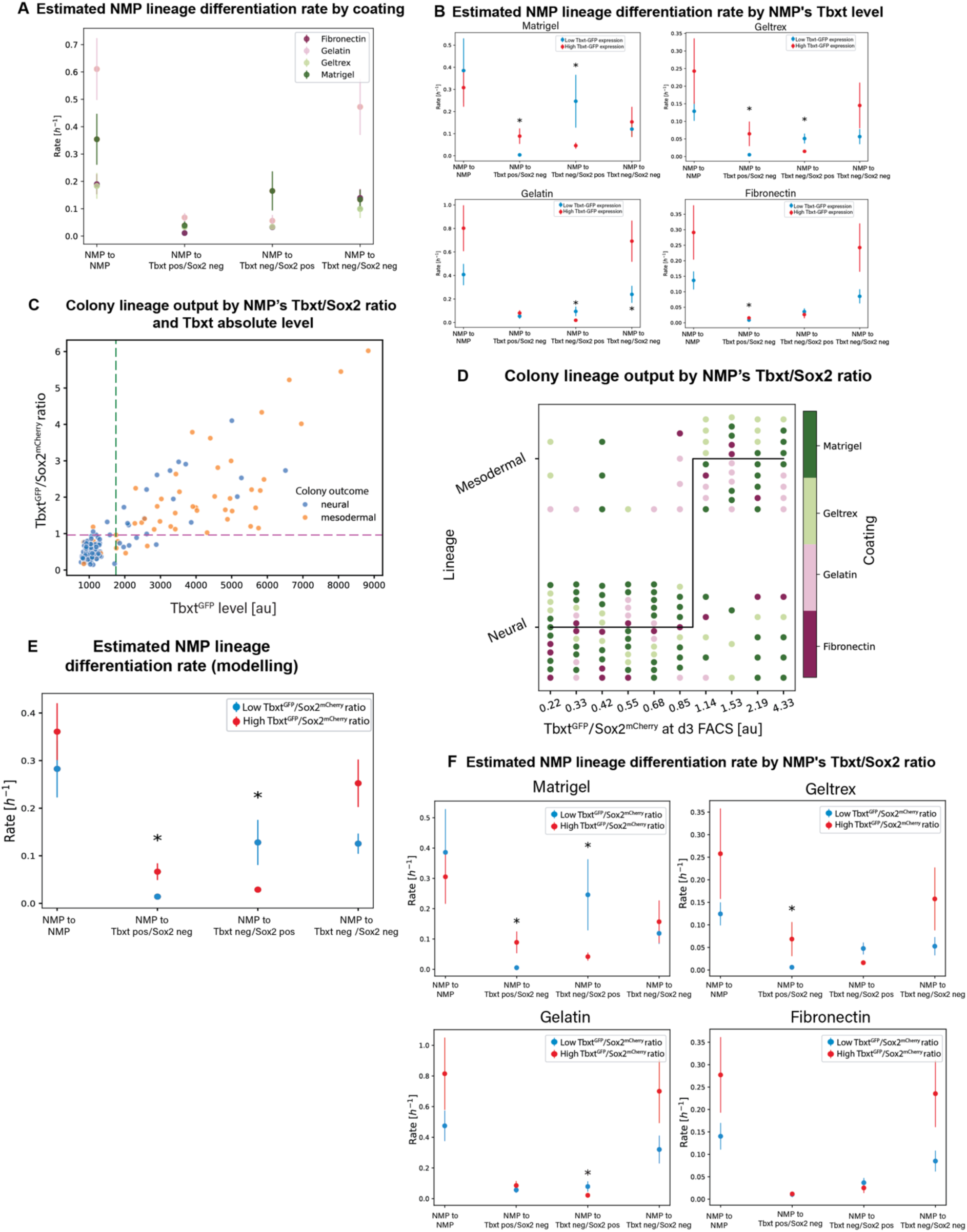
Quantitative analysis and mathematical modelling of clonal data using the Tbxt absolute levels Tbxt/Sox2 levels ratio. **A.** Lineage differentiation rates estimated with our mathematical model (Tbxt absolute levels), coloured by coating: matrigel (dark green), geltrex (light green), gelatin (pink) and fibronectin (fuchsia). Dots are the mean with error bars, the standard error of the mean**. B.** Differentation lineage rates estimated with our mathematical model (Tbxt absolute levels) sorted by coating used. The initial NMP cells were above (in red) or below (in blue) a TbxtGFP level threshold of 1745 arbitrary units (au) **C.** Colony lineage output when plotting Tbxt/Sox2 ratio over Tbxt absolute numbers. GFP/mCherry threshold value = 0.97 **D.** NMP lineage output as a function of initial Tbxt^GFP^ /Sox2^mCh^ ratio. Each dot represents one colony/experiment, with substrates used: matrigel (dark green), geltrex (light green), gelatin (pink) and fibronectin (fuchsia). The ratio reported is the middle value in the bin. **E.** Mathematical model to estimate fate in index sorted NMPs. The rate indicates whether single founder NMPs were above (red) or below (blue) the initial Tbxt^GFP^ /Sox2^mCh^ threshold ratio (0.97), as a function of their colony output at the end of the experiment. The dots and the error bars correspond to the mean and the standard error of the mean, respectively. Asterisks indicate significant differences obtained after applying a Mann-Whitney statistical test (p < 0.05). **F.** Differentiation rates of individual lineages estimated with our mathematical model sorted by coating used. The initial NMP cells were above (in red) or below (in blue) the Tbxt^GFP^ /Sox2^mCh^ threshold ratio. Dots are the mean with error bars, the standard error of the mean. Asterisks indicate statistically significant differences (*p* < 0.05) obtained after applying a Mann-Whitney statistical test.

**Table S1.**
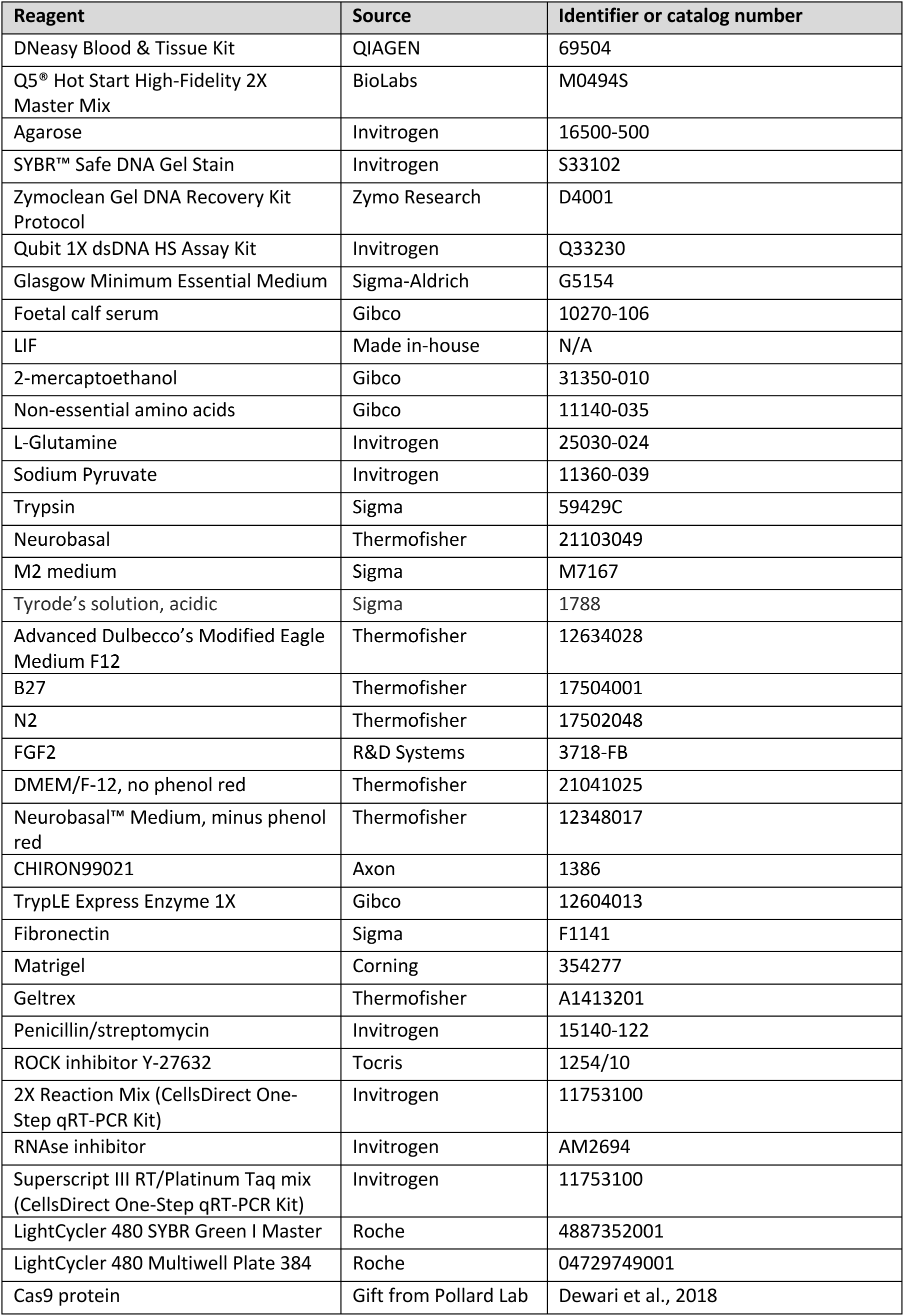
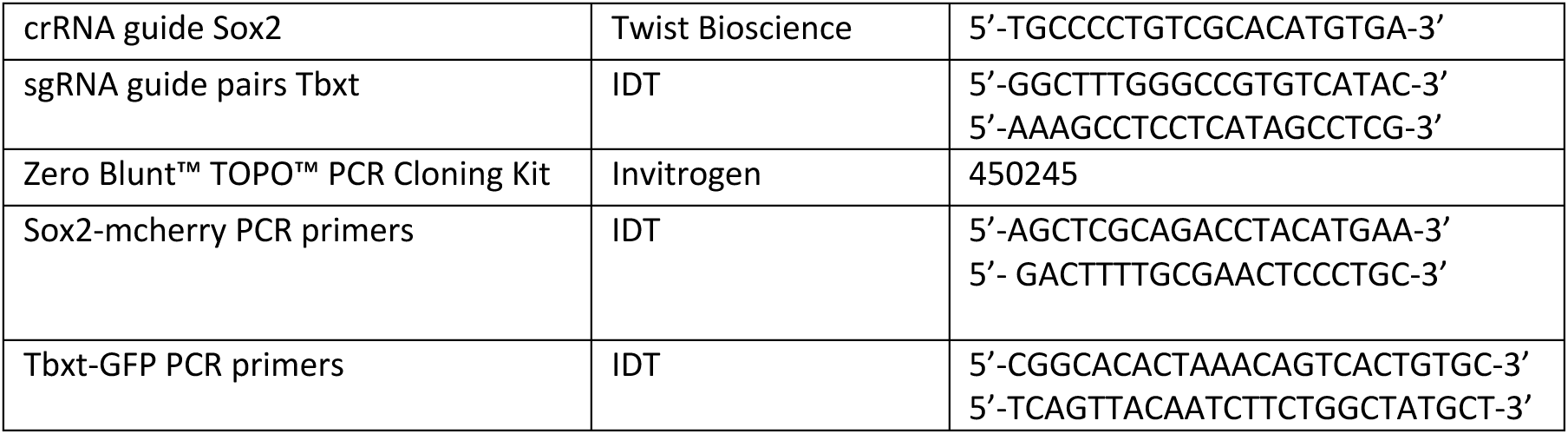

